# Neanderthal-derived variants shape craniofacial enhancer activity at a human disease locus

**DOI:** 10.1101/2024.09.24.614243

**Authors:** Kirsty Uttley, Hannah J Jüllig, Carlo De Angelis, Julia MT Auer, Ewa Ozga, Hemant Bengani, Hannah K Long

## Abstract

Facial appearance is one of the most variable morphological traits in humans, influenced by both rare and common genetic variants that can impact facial form between individuals and in disease. Deletion of an enhancer cluster 1.45 megabases upstream of the *SOX9* gene (EC1.45) results in Pierre Robin sequence, a human craniofacial disorder characterised by underdevelopment of the lower jaw and frequently associated with cleft palate. We reasoned that single nucleotide variants in EC1.45 may cause more subtle alterations to facial morphology. Here, we took advantage of recent human evolution, and the distinct morphology of the Neanderthal lower jaw, to investigate the impact of three Neanderthal-derived single nucleotide variants on EC1.45 function and jaw development. Utilising a dual enhancer-reporter system in zebrafish, we observed enhanced Neanderthal regulatory activity relative to the human orthologue during a specific developmental window. At this same stage, we show that EC1.45 appears to be selectively active in neural crest-derived progenitor cells which lie in close apposition with and are transcriptionally related to precartilaginous condensations that contribute to craniofacial skeletal development. To examine the potential consequences of increased *SOX9* expression in this specific cellular population during jaw development, we overexpressed human SOX9 specifically in EC1.45-active cells and observed an increase in the volume of developing cartilaginous precursors. Taken together, our work implicates Neanderthal-derived variants in increased regulatory activity for a disease-associated enhancer with the potential to impact craniofacial skeletal development and jaw morphology across recent hominin evolution.

## Introduction

Many human diseases implicate genetic changes in the non-coding genome, including both Mendelian (French and Edwards 2020; Spielmann, Lupiáñez, and Mundlos 2018; Zhang and Lupski 2015) and complex disease (Claringbould and Zaugg 2021). In many cases these genetic variations have been proposed to perturb gene regulatory sequences termed enhancers (Long, Prescott, and Wysocka 2016), leading to mis-regulation of target gene expression and perturbation of normal development (Claringbould and Zaugg 2021). An archetypal example are the structural variants at the *SOX9* locus in patients with Pierre Robin sequence (PRS) (Tan and Farlie 2013; Robin 1994). Unlike *SOX9* coding mutations which cause the severe congenital disorder campomelic dysplasia (characterised by bowed long limbs, disorders of sex determination, and craniofacial defects (Wagner et al. 1994)), PRS is a malformation characterised by underdevelopment of the lower jaw (micrognathia), a backwards displacement of the tongue (glossoptosis), airway obstruction, and frequent incidence of cleft palate (Amarillo, Dipple, and Quintero-Rivera 2013; Benko et al. 2009; C T Gordon et al. 2009; Christopher T. Gordon et al. 2014). Genetic alterations in PRS patients include large deletions and translocation breakpoints at distances over 1.2 Mb upstream of *SOX9* (Amarillo, Dipple, and Quintero-Rivera 2013; Benko et al. 2009; Christopher T. Gordon et al. 2014). We previously characterised two enhancer clusters that lie upstream of these translocation breakpoints and are ablated by at least one described PRS patient deletion (Long et al. 2020). These enhancer clusters were specifically active in *in vitro*-derived cranial neural crest cells (CNCCs), multipotent and transient cells that give rise to the majority of craniofacial structures (Bronner and LeDouarin 2012), and were decommissioned during differentiation to chondrocytes (cartilage-forming cells) (Long et al. 2020). The strongest and most distal of the enhancer clusters, 1.45 Mb upstream of *SOX9* (EC1.45) (Figure 1A), exhibited activity in the developing mouse craniofacial region and limb bud. We further demonstrated that genetic ablation of the orthologous mouse element impacted lower jaw development and lead to reduced postnatal fitness, emphasising a key role for this regulatory element in shaping jaw morphology and function (Long et al. 2020). While ablation of the EC1.45 region in human patients is associated with severe underdevelopment of the lower jaw, it is likely that sequence variation within the EC1.45 element may also alter enhancer activity, impacting *SOX9* developmental expression and ultimately shaping jaw morphology.

**Figure 1.**
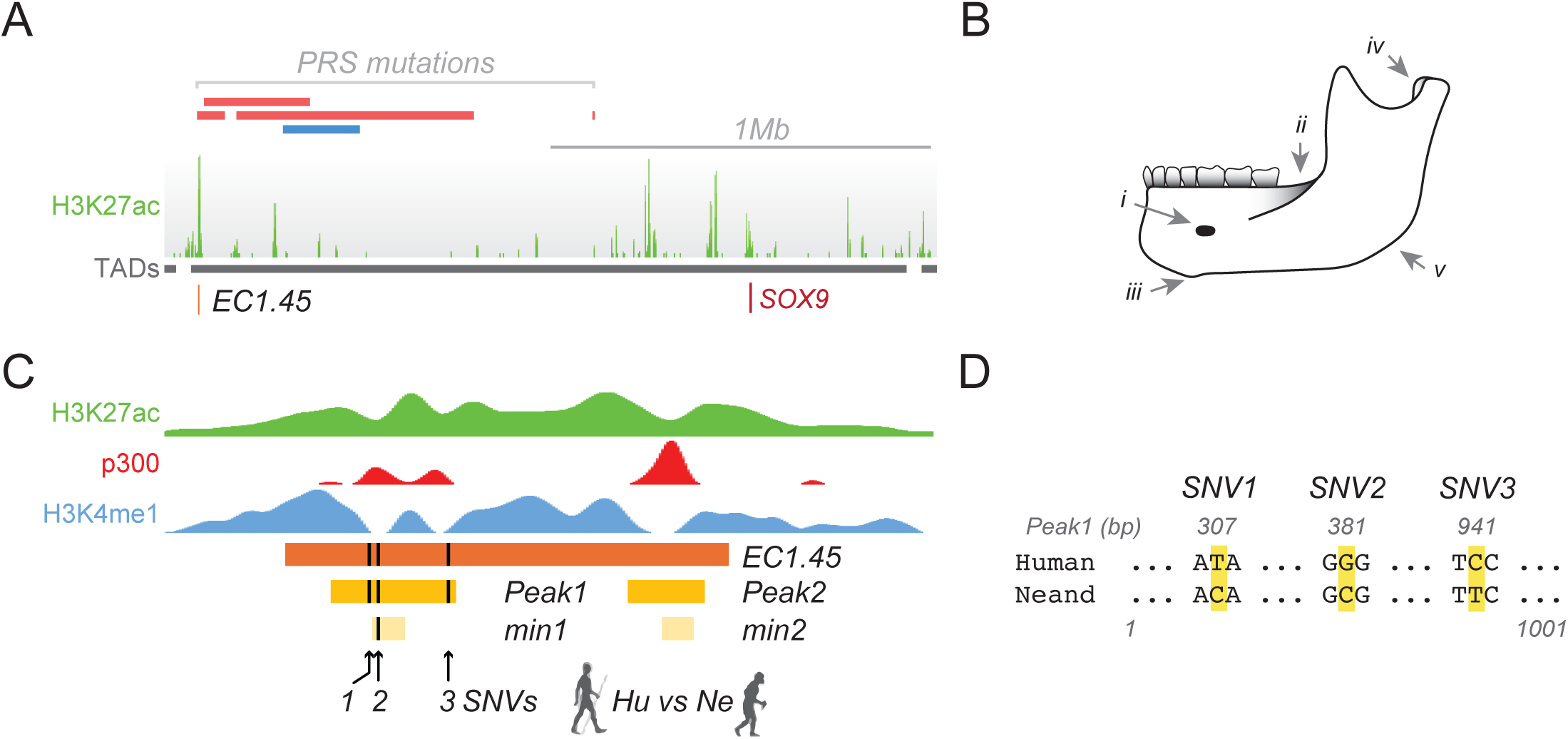
Three Neanderthal-derived single nucleotide variants overlap a *SOX9* disease- associated enhancer cluster. (A) The cranial neural crest cell-specific enhancer cluster EC1.45 falls within a deletion hotspot and upstream of a cluster of translocation breakpoints identified in patients with Pierre Robin sequence. H3K27ac ChIP-seq from CNCCs (Prescott et al. 2015) (green), topological associated domains (TADs) (grey) (Dixon et al. 2012), PRS patient deletions (red) and a cluster of translocation breakpoints (blue) (Amarillo, Dipple, and Quintero-Rivera 2013; Benko et al. 2009; C T Gordon et al. 2009; Christopher T. Gordon et al. 2014). (B) The Neanderthal jaw exhibits several derived morphological features (apomorphies), including i) posterior position of mental foramen, ii) retromolar gap, iii) posterior positioning of anterior marginal tubercle, iv) laterally expanded condyle, and v) truncated gonion. Adapted from (Rosas 2001). (C) Schematic of the EC1.45 region, highlighting Peak1-2 and min1-2 (Long et al. 2020). Three SNVs fall within the EC1.45 enhancer cluster, all within the Peak1 region, one of which overlaps the min1 region. ChIP-seq for H3K27ac (green), p300 (red) and H3K4me1 (blue) highlight the enhancer-associated chromatin features of the locus in CNCCs. Data from (Prescott et al. 2015). (D) DNA sequence alignment highlighting the sequence context for the 3 Neanderthal SNVs within the EC1.45 Peak1 region compared to human.

The morphology of the lower jaw is highly variable among vertebrate species (Woronowicz and Schneider 2019), associated with diversity in diet, feeding methods, vocalisation, communication, and gait (Coombs et al. 2024; Morales-García et al. 2021). This morphological diversity of jaw structure is exemplified by the striking radiation observed across Darwin’s finches and cichlid fishes (Abzhanov et al. 2004; Albertson and Kocher 2006). The fossil record also shows dramatic changes in skeletal form across an estimated 4 million years of hominin evolution (Bergmann et al. 2021; Lacruz et al. 2019). Hominins have had exceptionally rapid rates of mandibular shape evolution compared to other primate clades, hypothesised to be influenced by changes in diet and reduction in size of canine teeth (Bergmann et al. 2021; Raia et al. 2018). Lower jaw formation is initiated through condensation of CNCC-derived mesenchymal cells within pharyngeal arch 1 (PA1) to form precartilaginous condensations (PCCs). Cells within the condensing mesenchyme mature and differentiate to chondrocytes that form Meckel’s cartilage (Svandova et al. 2020; Gillis 2019). PCCs form within a wider zone of mesenchymal progenitor cells, and cells which do not contribute directly to the condensation may form perichondral cells on the outside of the cartilage, or intramembranous bone, tendon or ligament (Paudel et al. 2022; Hall 2015). Once established, the Meckel’s cartilage template is proposed to act as a model for the adult mandible, with the mandibular bone ossifying alongside and encasing the intermediate portion of Meckel’s cartilage which eventually is then eventually resorbed or undergoes transdifferentiation to other tissue-types (Svandova et al. 2020). Therefore, alteration to neural crest development, PCC patterning, cartilage or bone formation could each impact the ultimate shape and size of the mandible (Pitirri et al. 2022).

Sequencing of ancient hominin and other primate genomes has revealed that coding sequences tend to be highly conserved across the timescales separating modern humans from our close hominin relatives, including the Neanderthals that are predicted to have diverged around 500,000 years ago (Suntsova and Buzdin 2020; Weiss et al. 2021). By contrast, non-coding sequence changes are much more abundant (Yan and McCoy 2020). Unlike coding variation which is likely to have a pleiotropic impact on gene activity in all tissues, DNA sequence changes in non-coding sequences such as enhancers can drive altered morphology by modulating gene expression in a cell type specific and spatiotemporal manner (Long, Prescott, and Wysocka 2016; Prescott et al. 2015). Indeed, across evolutionary timescales, there are many examples of morphological change driven by alterations in developmental gene expression patterns (Frankel et al. 2011; Long, Prescott, and Wysocka 2016; Rubinstein and De Souza 2013).

We previously observed that EC1.45 overlaps with a differentially methylated region (DMR) that is hypomethylated uniquely in Neanderthal bone samples, (Gokhman et al. 2014; 2020) suggestive of a gain of function variant in the Neanderthal enhancer (Long et al. 2020). Here, we explored the potential impact of three Neanderthal-derived single nucleotide variants (SNVs) within EC1.45 on enhancer function and jaw morphological variation using zebrafish enhancer-reporter lines. Zebrafish represents a powerful model system for exploring alterations in hominin gene regulatory mechanisms due to broadly conserved craniofacial gene regulatory networks, well-characterised and orthologous craniofacial development to human (including formation of Meckel’s cartilage), external and transparent development, and rapid generation times (Mork and Crump 2015; Raterman et al. 2020). *sox9a* mutant embryos also exhibit craniofacial and cartilage developmental defects analogous to human campomelic dysplasia patients, with more severe phenotypes observed for *sox9a/sox9b* double mutant embryos, highlighting the conservation of SOX9 function in vertebrate facial formation (Yan et al. 2002; 2005).

Here, we first characterised the spatiotemporal developmental expression patterns for the human EC1.45 enhancer cluster during zebrafish development. We identified enhancer activity in a population of neural crest progenitor cell populations that are in proximity to and are transcriptionally related to precartilaginous condensations (PCCs) that give rise to parts of the craniofacial skeleton including aspects of the palate and Meckel’s cartilage. Utilising a dual enhancer-reporter strategy, we revealed that the Neanderthal orthologue of EC1.45 exhibits greater enhancer activity compared to human. Mimicking this increased activity, we demonstrated that overexpression of human SOX9 in EC1.45-active cells caused an increase in the size of the embryonic craniofacial precartilaginous template, indicating a possible link to jaw morphological changes observed between humans and Neanderthals. Our data implicate ancient hominin sequence variants in driving differential enhancer activity in an intriguing population of craniofacial progenitors and provide functional insights into the impact of increased SOX9 expression in these cells. Ultimately this work explores gene regulatory mechanisms that were altered in the Neanderthal lineage and may have contributed to an increase in SOX9 expression during development and a subsequent alteration to jaw morphology.

## Results

### Neanderthal-specific variants in a human disease-associated enhancer cluster

Given the pathogenic consequences associated with ablation of the EC1.45 enhancer cluster, we hypothesised that DNA sequence changes in this regulatory element may cause more subtle, normal-range morphological change in the human population, or altered morphology across evolutionary time (Figure 1A). To explore this possibility, we considered recent human evolution, and the distinct jaw morphological features between anatomically modern humans and Neanderthals. From the fossil record Neanderthal-specific jaw features, or evolutionary novelties (apomorphies), have been described including a retromolar space, a truncated gonion, and a posterior position of the mental foramen (Figure 1B) (Bergmann et al. 2021; Rosas 2001). From three high quality genomes available for Neanderthal we identified three SNVs overlapping EC1.45 (Figure 1C) (Prüfer et al. 2014; Green et al. 2010; Mafessoni et al. 2020). Two of the Neanderthal variants are present in all three high quality genomes, while the third (SNV3) is present only in one suggesting that it may be a sequencing artefact or have been polymorphic in the Neanderthal population (Figure 1D).

The EC1.45 element is comprised of two regions defined by p300 peaks from chromatin immunoprecipitation sequencing (ChIP-seq) called Peak1 and Peak2, that we previously narrowed down to two minimally-active sequences (min1 and min2) that recapitulate the regulatory activity of both EC1.45 and Peak1-2 by *in vitro* luciferase reporter assay (Long et al. 2020) (Figure 1C). One of the variants, SNV2, overlaps with the min1 region, potentially indicating a greater chance of functional impact. Furthermore, SNV2 creates a new CpG in the Neanderthal genome (Figure 1D), perhaps related to the alteration in CpG methylation status detected in ancient Neanderthal bone samples (Gokhman et al. 2014; 2020). Sequence variation is also observed for other non-human primate species in Peak1, but this is non-overlapping with the Neanderthal variants described here, and the Neanderthal variants are not present in gnomAD v4.0, apart from one instance of the potentially polymorphic SNV3 variant (1/152,152 rs563260668) (Karczewski et al. 2020). Notably, the three variants do not match the predicted ancestral state and therefore appear to be derived in Neanderthal, and thus may be associated with the appearance of Neanderthal-specific jaw features or apomorphies.

### EC1.45 enhancer activity is detected in cells directly adjacent to the developing craniofacial skeleton

To explore enhancer activity dynamics of human EC1.45 across development, we generated a zebrafish Tol2-mediated transgenic reporter line with the Peak1-2 region cloned upstream of the minimal gata2 promoter and eGFP (Figure 2A), which we named *Tg(HuEC1.45-P1P2:eGFP*), abbreviated to *Tg(HuP1P2:GFP)*. The expression pattern of the human EC1.45 Peak1-2 enhancer cluster was assessed by confocal imaging of *Tg(HuP1P2:GFP)* embryos crossed to the reporter line *Tg(sox10:mRFP)* (Kucenas et al. 2008; Kirby et al. 2006) (mRFP: membrane-bound red fluorescent protein). During zebrafish development, *sox10* is expressed in CNCCs, and continues to be expressed in the cranial PCCs which will form skeletal elements of the viscerocranium that includes the lower jaw, and the ethmoid plate of the neurocranium that forms the larval upper jaw (Kucenas et al. 2008; Mork and Crump 2015). At 1 day post fertilisation (dpf), eGFP was detected broadly in the frontonasal region (Supplementary Figure 1A). By 2 dpf, eGFP expression persisted in the frontonasal region, and appeared in a paired location adjacent to the mouth (Figure 2B and Supplementary Movies 1 and 2). Specifically, enhancer activity was detected alongside Meckel’s precartilaginous condensations and extended anteriorly along the oral cavity (Eames et al. 2013) (see Figure 2C and Supplementary Figure 1B for schematics of developing cartilage structures and relative location of EC1.45-P1P2 reporter activity). At 3 and 4 dpf, enhancer activity continued to be detectable adjacent to Meckel’s cartilage, at a lateral and slightly dorsal position, and appeared to extend into the Meckel’s precartilaginous template at the jaw joint region (Figure 2B-C, Supplementary Figure 1B and Supplementary Movie 3). Time-lapse imaging from 2-4 dpf confirmed that the paired Meckel’s-adjacent signal was the same population at 2, 3 and 4 dpf (Supplementary Movie 4). For the enhancer-positive cells in the frontonasal region, a posterior subset of these cells appeared to contribute to the forming palate between 2- 3 dpf, with eGFP-expression observed in the ethmoid plate at these stages (Supplementary Movie 5, arrow). This is consistent with previous work which used fate mapping to show that frontonasal CNCCs adjacent to the nasal epithelium populate the ethmoid plate (Swartz et al. 2011). Similar domains of enhancer activity were observed for *Tg(HuP1P2:GFP)* crossed to a *col2a1a* reporter line, *Tg(col2a1a:RFP)*, which also marks precartilaginous condensations and chondrocytes during craniofacial development (Dale and Topczewski 2011; Paudel et al. 2022) (Supplementary Figure 1B-C).

**Figure 2.**
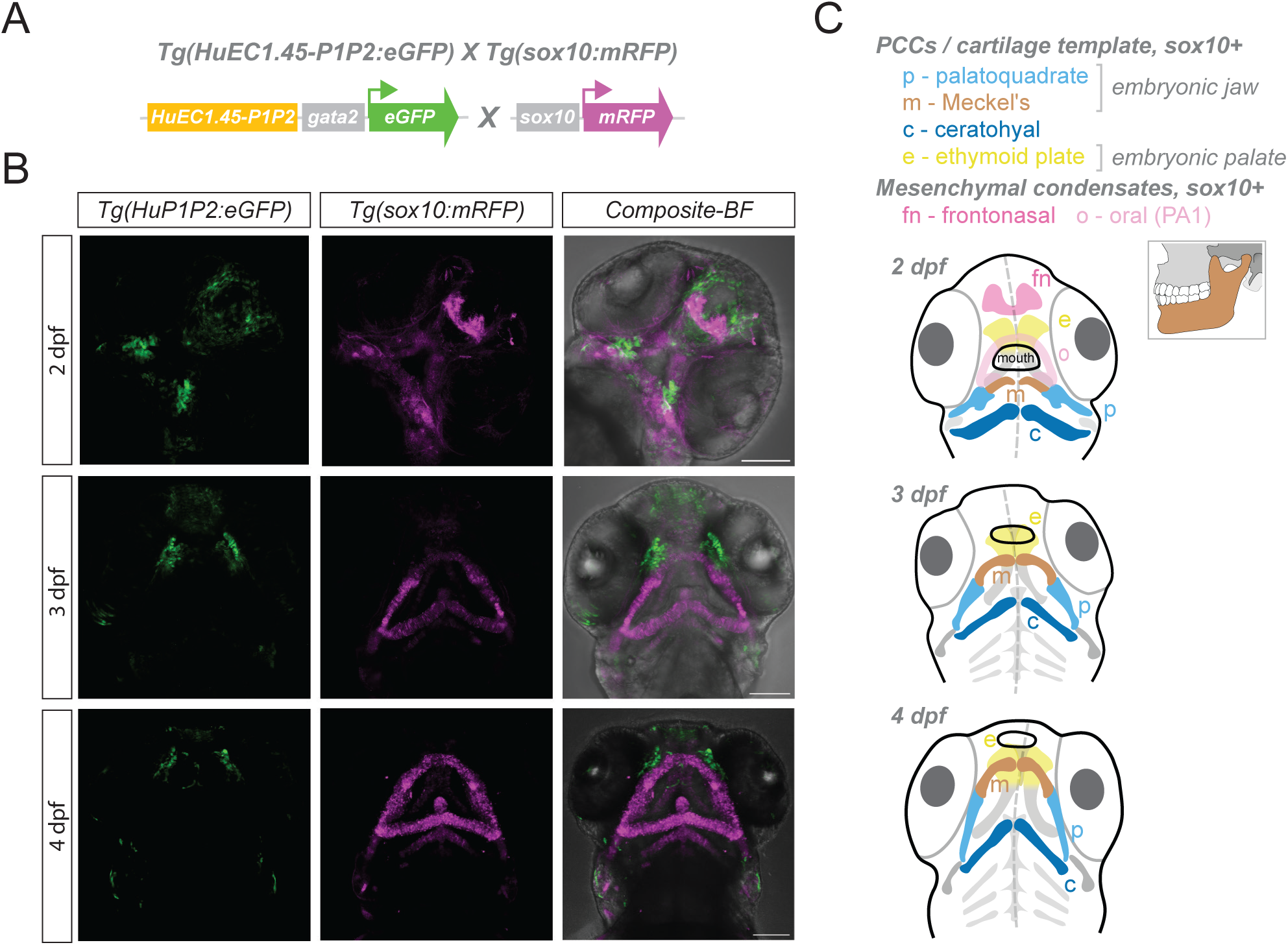
EC1.45 Peak1-2 is active during embryonic facial development in the frontonasal region and adjacent to the developing jaw. (A) Schematic of the cross between two transgenic lines, human EC1.45 Peak1-2 enhancer reporter *Tg(HuEC1.45-P1P2:eGFP)* and the *Tg(sox10:mRFP)* reporter. The Peak1-2 region is upstream of a minimal gata2 promoter driving eGFP expression and flanked by insulators and Tol2 sites. (B) Confocal microscopy images of the cranial region of embryos at 2, 3 and 4 dpf for *Tg(HuEC1.45-P1P2:eGFP)* crossed with *Tg(sox10:mRFP)*. Maximum intensity projections are shown for representative embryos. Scale bars 100 µm. (C) Schematics of zebrafish embryonic cranial region (ventral view) indicating the location of *sox10:mRFP* reporter activity in precartilaginous condensates (PCCs) which mature to cartilage templates from 2-4 dpf. Additional neural crest-derived mesenchymal populations are indicated at 2 dpf (fn and o). m – Meckel’s (corresponding to human lower jaw); p – palatoquadrate; e – ethmoid plate; c – ceratohyal; fn – frontonasal; o – oral; PA1 – pharyngeal arch 1.

The expression domains for HuEC1.45-P1P2 observed during zebrafish development were reminiscent of enhancer reporter activity we previously described for the mouse embryo (Long et al. 2020). At mouse embryonic day 9.5 (E9.5), EC1.45 was active in the developing frontonasal prominence, and at E11.5 in the lateral nasal process, medial nasal process, maxillary process, mandibular process, periocular mesenchyme, and limb bud. Strikingly, we also detected eGFP expression in the developing fin at 3 and 4 dpf for the *Tg(HuP1P2:GFP)* line (Supplementary Figure 1D), in keeping with the limb expression patterns observed during mouse development (Long et al. 2020) . Together, a zebrafish reporter of human EC1.45 regulatory activity matches key expression domains observed from mammalian development and provides increased temporal and spatial insights into the developmental activity of this disease-associated regulatory element, especially for a population of cells in proximity to the developing lower jaw.

### Neanderthal variants increase EC1.45 enhancer activity compared to human during early craniofacial development

To explore how the three Neanderthal-derived SNVs impact the activity of the EC1.45 enhancer cluster, we leveraged the Q-STARZ assay (Quantitative Spatial and Temporal Assessment of Regulatory element activity in Zebrafish) which enables the activity of two enhancers to be tested using a single enhancer reporter construct during zebrafish embryonic development (Figure 3A) (Bhatia et al. 2021; Uttley et al. 2023). Firstly, the Neanderthal Peak1-2 sequence was placed upstream of a minimal promoter and eGFP and separated by an insulator sequence from the human version of Peak1-2 driving expression of mCherry *Tg(NeP1P2:eGFP;HuP1P2:mCh)*, abbreviated to *Tg(Ne:GFP;Hu:Ch)* (Figure 3A). Stable transgenic lines were generated by Tol2- mediated transposition, and embryos were collected for imaging by confocal microscopy across the first 3 days of development. The spatial activity of Neanderthal Peak1-2 (eGFP signal) appeared highly similar to that of the human enhancer (mCherry signal) across development (Figure 3B and Supplementary Figure 2A-B). However, we observed that the absolute expression level of eGFP driven by Neanderthal Peak1-2 appeared to be detectably higher than for mCherry driven by human Peak1-2, especially at 2 dpf (Figure 3B). We recapitulated these results, with observed higher activity of the Neanderthal enhancer at 2 dpf, in a ‘swap’ line with the human enhancer upstream of eGFP, and the Neanderthal enhancer upstream of mCherry (*Tg(HuP1P2:eGFP;NeP1P2:mCh)*, abbreviated to *Tg(Hu:GFP;Ne:Ch)*) (Figure 3A-B). Finally, we created a transgenic line in which the expression of both eGFP and mCherry are controlled by the human enhancer sequence (*Tg(HuP1P2:eGFP;HuP1P2:mCh)*, abbreviated to *Tg(Hu:GFP;Hu:Ch)*) and as expected we did not observe a difference in eGFP and mCherry expression levels at these developmental stages (Figure 3A-B and Supplementary Figure 2A-B). To confirm these observations, we performed quantification of the mean fluorescence intensity for eGFP and mCherry in the frontonasal regions (including some cells contributing to the developing palate) and mandible-adjacent structures from 1-3 dpf (see Supplementary Figure 2C for representative images of surfaces generated to quantify expression). Confirming our observations, we detected significant differences in the mean fluorescence intensity for eGFP and mCherry at 2 dpf in the *Tg(Ne:GFP;Hu:Ch)* and *Tg(Hu:GFP;Ne:Ch)* lines, but not for the *Tg(Hu:GFP;Hu:Ch)* line (Figure 3C). Specifically, the Neanderthal enhancer drove significantly higher expression levels at this developmental stage for both the frontonasal and future jaw regions (Figure 3B-C). While we observe some evidence for a similar bias at 1 and 3 dpf, this was not statistically significant for the two swap lines (Supplementary Figure 2D-E). We therefore concluded that the three Neanderthal SNVs cause an increase in enhancer activity during zebrafish embryonic development in a temporally controlled manner, with the greatest impact detected at 2 dpf.

**Figure 3.**
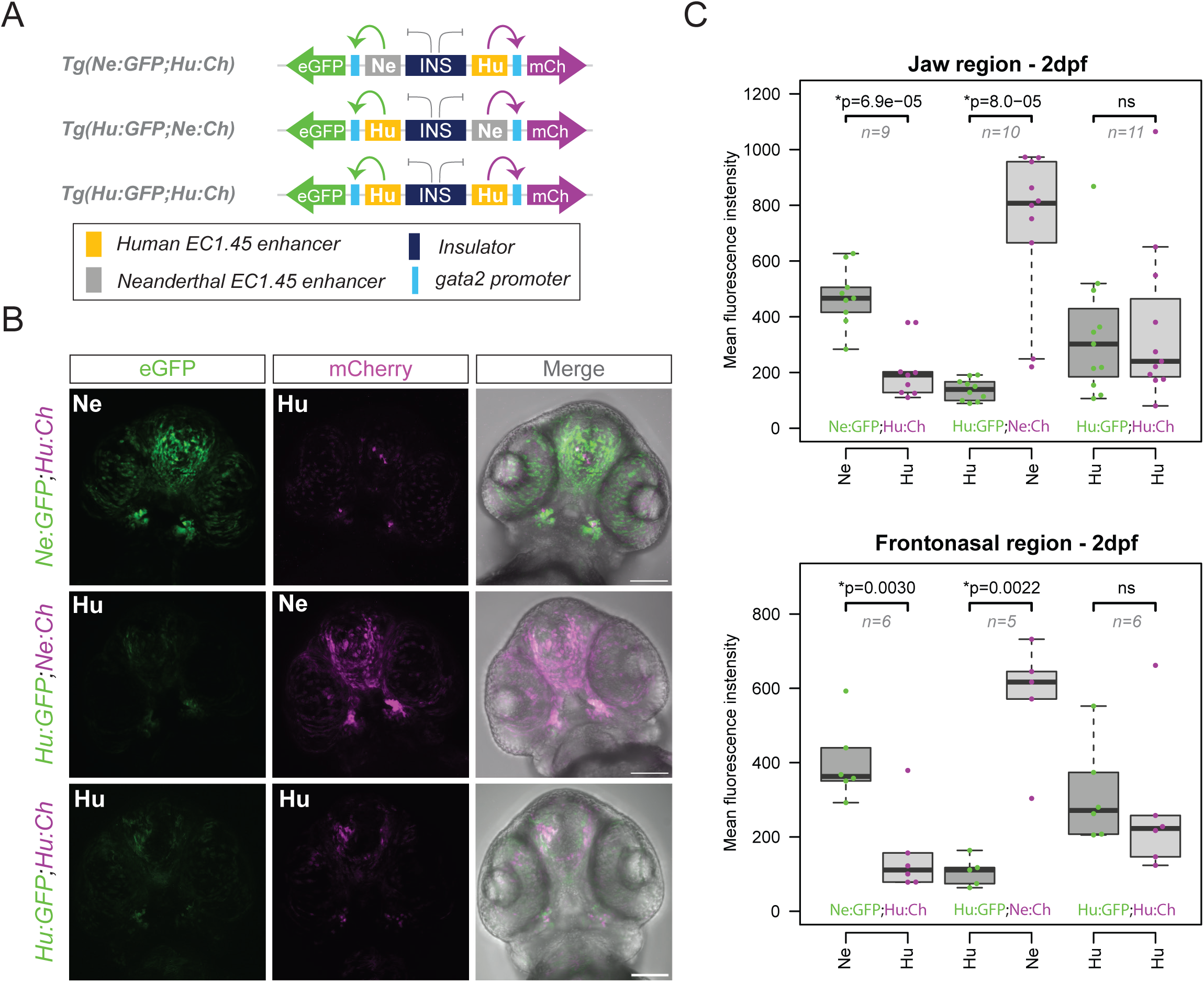
Neanderthal SNVs drive increased developmental enhancer activity compared to human. (A) Schematic of three Q-STARZ dual transgenic reporter lines created to compare human and Neanderthal EC1.45 enhancer element activity. For reporter line *Tg(Ne:GFP;Hu:Ch)*, the Neanderthal Peak1-2 sequence is upstream of eGFP, and human Peak1-2 upstream of mCherry. For reporter line *Tg(Hu:GFP;Ne:Ch)*, the enhancers are exchanged. And for reporter line *Tg(Hu:GFP;Hu:Ch)*, human Peak1-2 is upstream of both eGFP and mCherry. (B) Representative confocal images (maximum intensity projections) for embryos at 2 dpf, ventral view of cranial region. eGFP expression was observed to be greater than mCherry for the *Tg(Ne:GFP;Hu:Ch)* line, while mCherry expression was greater in the swap line compared to eGFP, *Tg(Hu:GFP;Ne:Ch)*. No difference was observed between fluorophores for the *Tg(Hu:GFP;Hu:Ch)* line. Scale bars 100 µm. (C) Quantification from (B) at 2 dpf of the jaw region (upper) and frontonasal region (lower). P values from a Wilcoxon signed-rank test are shown. See Supplementary Figure 2C for representative segmented regions used for quantification.

### EC1.45 is active in cranial neural crest and precartilaginous mesenchymal progenitor cells

To explore the relative activity and cell type specificity of the human versus Neanderthal EC1.45 Peak1-2 enhancers further, we performed single cell-RNA sequencing (scRNA-seq) for the two human-Neanderthal enhancer reporter lines at 2 dpf. Fluorescence-activated cell sorting (FACS) was used to isolate all cells that were eGFP-positive, mCherry-positive or double-positive from dissected embryonic cranial regions at 2 dpf (Figure 4A-B) This resulted in 40,000 cells for the *Tg(Hu:GFP;Ne:Ch)* line and 10,000 cells for the *Tg(Ne:GFP;Hu:Ch)* line which were processed as two separate samples (Figure 4B and Supplementary Figure 3A).

**Figure 4.**
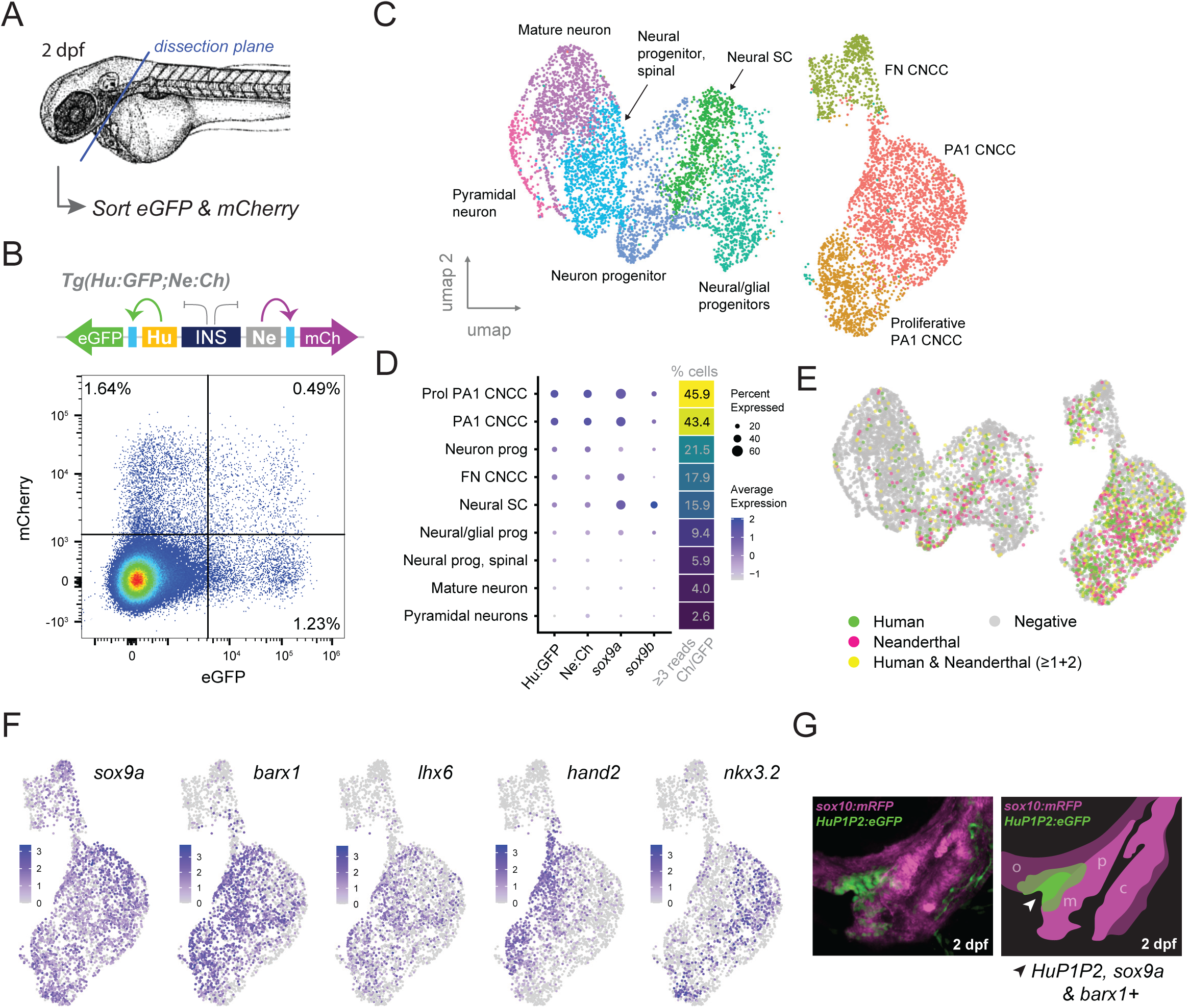
EC1.45 active cells transcriptionally resemble cranial neural crest cell-derived facial mesenchymal condensations. (A) Schematic illustrating dissection plane for isolating the cranial region from 2 dpf embryos for FACS followed by scRNA-seq. (B) FACS plot for cranial region cells dissected from the *Tg(Hu:GFP;Ne:Ch)* transgenic line. (C) Cells from *Tg(Hu:GFP;Ne:Ch)* embryos at 2 dpf, visualised on a Uniform Manifold Approximation and Projection (UMAP) plot created by Louvain clustering using Seurat. Clusters were annotated as certain cell types aided by marker gene expression and the Daniocell atlas. (D) Dot plot showing average expression level and percentage of cells expressing *eGFP*, *mCherry*, *sox9a* or *sox9b* (clusters from C). Percentage of cells which express greater than 3 reads of eGFP or mCherry are shown for each cluster (as for C). (E) Expression of eGFP (≥3 reads, green), mCherry (≥3 reads, magenta) or both (≥3 reads in total for eGFP and mCherry, yellow) in single cells visualised on UMAP plots (clustered as in C). (F) Expression of facial condensation marker genes (*sox9a*, *barx1* and *lhx6*) and genes which pattern pharyngeal arch 1 (*hand2* and *nkx3.2*) in single cells visualised on UMAP plots (subset of clusters from C). (G) Lateral image of developing craniofacial structures at 2 dpf for cross between *Tg(HuEC1.45- P1P2:eGFP)* and *Tg(sox10:mRFP)*. Schematic and arrowhead highlight the domain of activity for HuEC1.45-P1P2. m – Meckel’s PCC; p – palatoquadrate PCC; c – ceratohyal PCC; o – oral mesenchymal condensations.

Following quality control and filtering, we carried out K-nearest neighbour analysis and clustering on 8,467 cells from the *Tg(Hu:GFP;Ne:Ch)* line, yielding 9 distinct clusters. Cell type annotation for these clusters was performed through identification of cluster-specific marker genes with aid from the Daniocell atlas (Sur et al. 2023; Farrell et al. 2018) (Figure 4C, Supplementary Figure 3B and Supplementary Table 1). This annotation revealed three CNCC-like clusters marked by canonical neural crest marker genes such as *twist1a* and *snai1a/2*, including a frontonasal population (expressing *alx1* and *alx4a/b* (Mitchell et al. 2021)), and two clusters resembling pharyngeal arch 1 (PA1) populations (expressing *dlx* genes and *barx1* (Talbot, Johnson, and Kimmel 2010; Simões-Costa and Bronner 2015)), one of which appeared more proliferative (Figure 4C and Supplementary Figure 3B-C). A number of additional clusters were annotated as various neuronal cell types (Figure 4C and Supplementary Figure 3B). To identify enhancer-active cells within this dataset we plotted the expression of either eGFP or mCherry, finding a notable enrichment in the CNCC clusters, particularly those of PA1 (Figure 4D and Supplementary Figure 3D). Cells were then categorised as having enhancer activity for the human (eGFP-positive), Neanderthal (mCherry-positive), or both reporters (double-positive) by the presence of 3 or more eGFP and/or mCherry reads. Once again, the highest proportion of cells with three or more mCherry or eGFP reads were the two PA1 CNCC clusters (43.4 and 45.9%) suggesting that these cells exhibit the greatest enhancer activity at this stage (Figure 4D-E and Supplementary Table 2). In keeping with endogenous EC1.45 regulating human *SOX9* during development, enrichment of eGFP and mCherry expression in the CNCC clusters was concurrent with greater levels of *sox9a* expression (Figure 4D-E and Supplementary Figure 3D). The lower level of *sox9b* expression in these clusters is consistent with a lesser role for *sox9b* in craniofacial development in the zebrafish compared to *sox9a* (Supplementary Figure 3D) (Yan et al. 2005). In accordance with our imaging data, we did not observe any differences in the cell type identity of human versus Neanderthal active cells, which appear to broadly group together in the CNCC clusters (Figure 4D-E). Similar results were observed from the smaller sample of cells obtained from the *Tg(Ne:GFP;Hu:Ch)* line, where 407 cells were grouped into three clusters of PA1 CNCCs, FN CNCCs, and neuronal cells (Supplementary Figure 3E-F), where the eGFP and mCherry expression was again most enriched in the PA1 CNCC cluster (Supplementary Figure 3G).

To further examine the neural crest cell clusters with highest enhancer activity, we investigated genes known to pattern PA1 at this stage of development. We observed expression of *barx1*, *lhx6*, *hand2* and *nkx3.2* in subsets of the neural crest cells from the two PA1 clusters (Figure 4F and Supplementary Figure 3C). Mirroring the known spatial expression of these markers, we found that *hand2* (expressed in the ventral domain of PA1) and *nkx3.2* (expressed in the joint forming intermediate domain of PA1) were expressed in a mutually exclusive and restricted pattern, while *barx1* and *lhx6* (expressed in both dorsal and ventral regions of PA1) exhibited more broad expression across the cluster, including an overlap with *hand2* (Figure 4F) (Nichols et al. 2013; Paudel et al. 2022). Comparing this to the pattern of eGFP and mCherry expression, EC1.45 Peak1-2 enhancer-active cells are broadly distributed across the PA1 clusters, similar to the *barx1* and *lhx6* expression patterns (Figure 4E-F). This is intriguing given published work showing a key role for *barx1* and *lhx6* in jaw mesenchymal condensations which mature to form PCCs and form facial cartilages (Nichols et al. 2013; Paudel et al. 2022). Significantly, *barx1-*expressing cells in the jaw region have been shown to activate *sox9a* during maturation into PCCs (Paudel et al. 2022). Furthermore, *barx1* mutant embryos exhibit reduced *sox9a* and *sox10* expression in the developing jaw, and defects in chondrogenesis (Nichols et al. 2013). The association of EC1.45 enhancer activity with mesenchymal condensation markers is compelling, as it supports our hypothesis from enhancer reporter imaging that EC1.45 Peak1-2 is active in the vicinity of Meckel’s PCCs from 2 dpf (Figure 2B-C, Figure 4G and Supplementary Figure 1B). Together, scRNA-seq from two dual enhancer reporter lines supports the conclusion that Neanderthal SNVs do not appear to impact the cell type-specific activity of the EC1.45 regulatory element during early zebrafish craniofacial development. Instead, Neanderthal regulatory variants appear to drive stronger reporter activity in the face, including in cells that appear primed to contribute to cartilage formation in the developing jaw region at 2 dpf.

To further explore which tissues EC1.45-positive cells give rise to, and whether we observe direct contribution to jaw cartilage formation, we created a Cre-driver construct controlled by the *HuEC1.45-P1P2* enhancer sequence (referred to as *HuEC1.45-P1P2:Cre)*. We decided to leverage a lineage-tracing strategy as our previous work had suggested that the EC1.45 enhancer is decommissioned during chondrogenesis and therefore the enhancer reporter signal was not expected to persist in the developing cartilage (Long et al. 2020). The *HuEC1.45-P1P2:Cre* construct was injected with Tol2 mRNA into 1-cell embryos resulting from a cross between a lineage tracing reporter line *Tg(ubi:loxP-AmCyan-loxP-ZsYellow)*, *Tg(ubi:CSY)* for short, and the *Tg(sox10:mRFP)* line (Supplementary Figure 4A). By imaging ZsYellow expression, this approach facilitated tracking the progeny of enhancer-active cells which had expressed the Cre driver with comparison to the *sox10:mRFP* reporter which marks neural crest and precartilaginous condensates. Injected embryos were screened at 1 dpf for craniofacial signal, and further screened at 2 dpf for signal in the paired Meckel’s adjacent region characteristic of *HuEC1.45- P1P2* enhancer activity (Supplementary Figure 4Bi). We identified embryos with ZsYellow signal in the expected tissues at these stages, along with some spurious recombination as is expected for F0 embryos (Supplementary Figure 4B). For one embryo which showed ZsYellow-positive cells adjacent to the Meckel’s PCC at 2 dpf, we carried out time-lapse imaging for 12 hours from 2 dpf (Supplementary Figure 4Bi and Supplementary Movie 6) and observed emergence of a ZsYellow-positive, *sox10*-positive cell overlapping with Meckel’s cartilage. ZsYellow-positive, *sox10*-positive cells were also detected in the developing palatoquadrate PCC (Supplementary Figure 4Bi). At later developmental stages we also identified embryos which exhibited ZsYellow signal in the expected enhancer pattern, in addition to ZsYellow expression in cells within the adjacent Meckel’s cartilage at 3 and 4 dpf, including in cells with characteristic elongated chondrocyte morphology (Supplementary Figure 4Bii-iii, arrowheads).

Based on these observations, we returned to our enhancer reporter line *Tg(HuP1P2:GFP)* to investigate whether this line could be used for short-term lineage tracing (Paudel et al. 2022). From our earlier *in vitro* studies, we expect EC1.45 enhancer to be turned off during chondrogenesis (Long et al, 2020), however the eGFP reporter may persist for longer than the enhancer is active and reporter transcription has ceased. Indeed, at 2 dpf we observed eGFP-positive cells overlapping with *sox10* expression in the Meckel’s PCC for the *Tg(HuP1P2:GFP)* line (Supplementary Figure 5A, white arrowheads, Figure 4G and Supplementary Movie 1), and we repeated this observation with a cross to the *Tg(col2a1a:RFP)* line (Supplementary Figure 5B, white arrowhead). To confirm we could also observe this activity in an independent enhancer reporter line, we crossed the *Tg(Ne:GFP;Hu:Ch)* line to the *sox10* reporter, and again observed eGFP-positive cells within the Meckel’s PCC marked by mRFP expression, this time driven by the Neanderthal enhancer (Supplementary Figure 5C, white arrowheads). Together these results support the model that the *EC1.45-P1P2* enhancer active cells can contribute to PCCs and give rise to mature chondrocytes of the developing lower jaw.

### SOX9 overexpression in EC1.45-positive lineage impacts precartilaginous mesenchymal condensations

Our observation that Neanderthal-specific DNA sequence alterations increase EC1.45 enhancer activity implies an associated increase in endogenous *SOX9* expression during embryonic facial development, potentially affecting jaw shape. We therefore reasoned we could mimic this change through transient overexpression of human SOX9 in EC1.45 active cells during zebrafish development, to reveal how increased enhancer activity may impact jaw morphology. We therefore created a Tol2 construct in which *HuEC1.45-P1P2* controls expression of human *SOX9*, followed by the ribosome skipping sequence T2A and eGFP, *HuP1P2:hSOX9-T2A-eGFP* (Figure 5A). We injected the overexpression construct together with Tol2 mRNA into 1-cell *Tg(sox10:mRFP)* embryos to visualise the impact on craniofacial skeletal development, using the *HuEC1.45-P1P2:eGFP* Tol2 construct as a control (Figure 5A).

**Figure 5.**
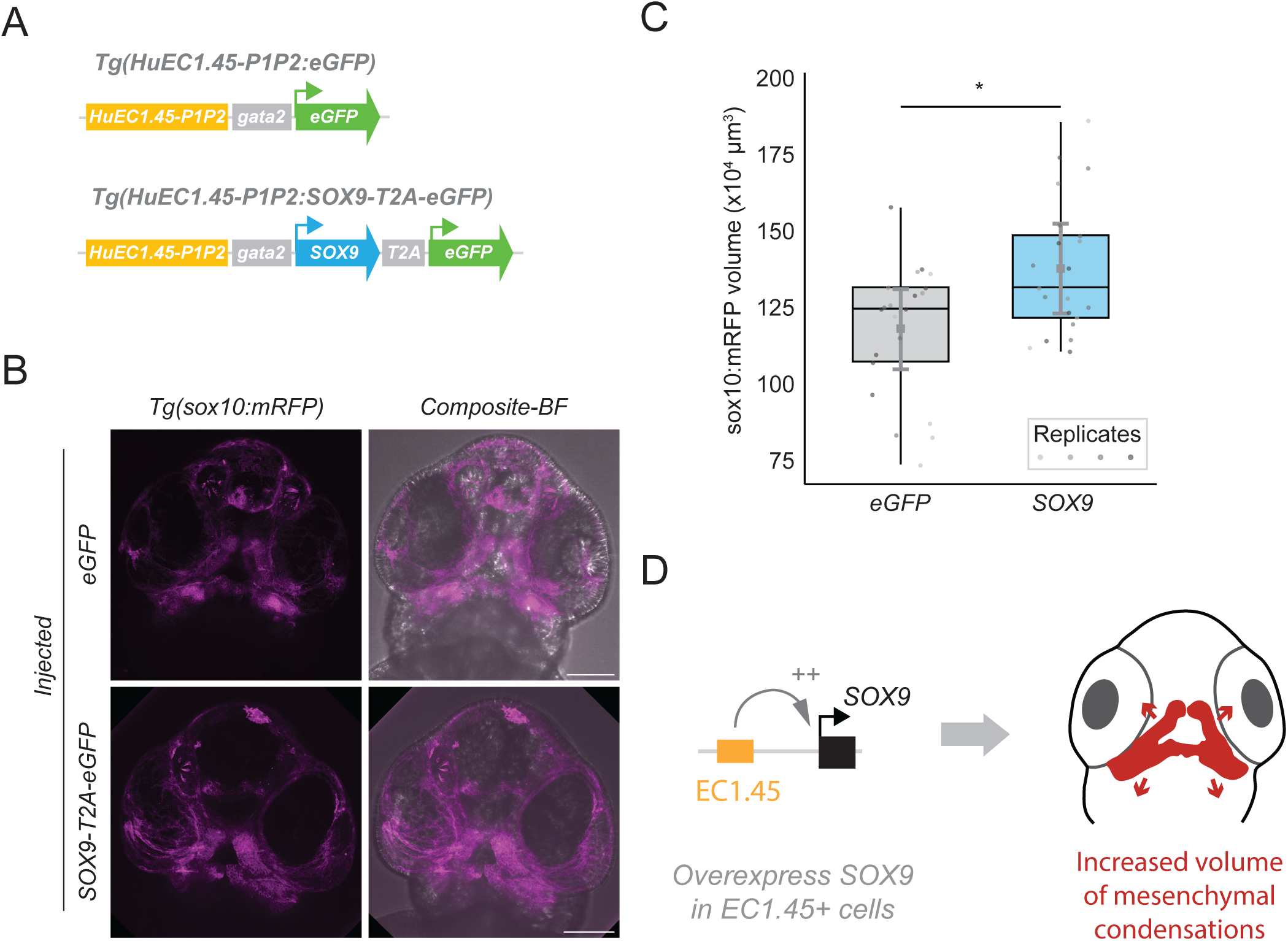
Overexpression of human SOX9 during zebrafish embryonic development in EC1.45 active cells impacts craniofacial development. (A) Schematic of the *HuEC1.45-P1P2:SOX9-T2A-eGFP* construct. The Peak1-2 region is upstream of a minimal gata2 promoter driving human SOX9 expression, followed by T2A-eGFP. The *HuEC1.45-P1P2:eGFP* construct from Figure 2A was used as a control. (B) Representative maximum intensity projections of confocal microscopy images for ventral view of 2 dpf embryonic craniofacial region. The *Tg(sox10:mRFP)* reporter was used to create a surface encompassing the mesenchymal cells of the Meckel’s and palatoquadrate precartilaginous condensations and the regions extending along the oral ectoderm (see Supplementary Figure 6C for representative segmented surfaces). Segmented volume from eGFP injected image = 126 x10^4^ µm^3^; SOX9-T2A-eGFP injected image = 133 x10^4^ µm^3^. (C) Quantification of *sox10:mRFP*-marked segmented volumes from (B) reveals that volume of *sox10:mRFP* positive region increases upon SOX9 overexpression compared to eGFP overexpression. Boxplots show the distribution of volumes between eGFP or SOX9 injected embryos at 2 dpf. For eGFP group mean = 120 x10^4^ µm^3^, median = 126 x10^4^ µm^3^; for SOX9 group mean = 139 x10^4^ µm^3^, median = 133 x10^4^ µm^3^. Grey error bars indicate the 95% credible intervals for the Bayesian mixed effects model posterior estimates, and the grey squares indicate the posterior mean estimates. The estimated increase in *sox10:mRFP*-positive volume for the SOX9 injected group was 19.6 x10^4^ µm^3^ (95% probability indicating a credible interval between 3.3 x10^4^ µm^3^ to 36.5 x10^4^ µm^3^). Wilcoxon test p=0.031. (D) Schematic depicting that overexpression of human SOX9 in EC1.45 active cells lead to an increased volume of mesenchymal condensations at 2 dpf.

Initially, injection of the SOX9 overexpression plasmid caused a spectrum of developmental phenotypes at 1 dpf. We therefore reduced the amount of SOX9 plasmid injected by combining with the eGFP-only construct at a 1:1 or 1:3 ratio. At these reduced SOX9 levels, the majority of injected embryos appeared grossly normal with milder and less frequent developmental defects (Supplementary Figure 6A). Only embryos with normal morphology were selected for imaging and analysis. For both *HuSOX9-T2A-eGFP* and *eGFP* overexpression we observed mosaic eGFP expression consistently in the face at 1 dpf, indicating success of the injections (Supplementary Figure 6B). Embryos from four replicate experiments with detectable eGFP expression and no overt developmental abnormalities were selected for confocal imaging at 2 dpf (Figure 5B). To assess the impact of SOX9 overexpression, we quantified the volume of the mRFP-positive mesenchymal condensates at the jaw region, including the Meckel’s PCC from the *Tg(sox10:mRFP)* injected transgenic embryos (Supplementary Figure 6C). Strikingly, this revealed a small but significant increase in the volume of the mRFP-positive region in the SOX9 overexpression embryos compared to eGFP overexpression alone (Figure 5C, Wilcoxon signed-rank test, p<0.05). We also performed a Bayesian mixed-effects model to control for multiple replicates and variation between injections. This model supported the finding that overexpression of SOX9 driven by the human EC1.45 Peak1-2 enhancer cluster significantly increases the mRFP-positive volume marking neural crest and PCCs (Figure 5C, posterior probability >95%). These results therefore suggest that overexpression of human SOX9 in cells where EC1.45 is active impacts the development of PCCs that will go on to form aspects of the craniofacial skeleton (Figure 5D).

In summary, we have uncovered an increase in activity for the Neanderthal EC1.45 orthologous enhancer cluster during craniofacial development and characterised regulatory activity in neural crest-derived mesenchymal cells, including in the jaw-forming region. These cells appeared transcriptionally related to a pre-cartilaginous state, associated with *barx1* expression, and by lineage tracing can contribute to PCC formation. We further demonstrated that overexpression of SOX9 in EC1.45-active cells can impact the volume of condensations associated with jaw formation. We therefore propose that alteration of the EC1.45 enhancer sequence during Neanderthal evolution may have promoted an increase in SOX9 expression during a window of craniofacial development that could contribute to altered abundance or morphology of cartilaginous precursors. Ultimately, while the Neanderthal SNVs in the EC1.45 enhancer cannot alone explain mandibular morphological variation between human and Neanderthal, these regulatory changes may have contributed to increased SOX9 expression during development leading to alteration of the craniofacial skeletal template and subtle morphological changes in jaw shape (Figure 6).

**Figure 6.**
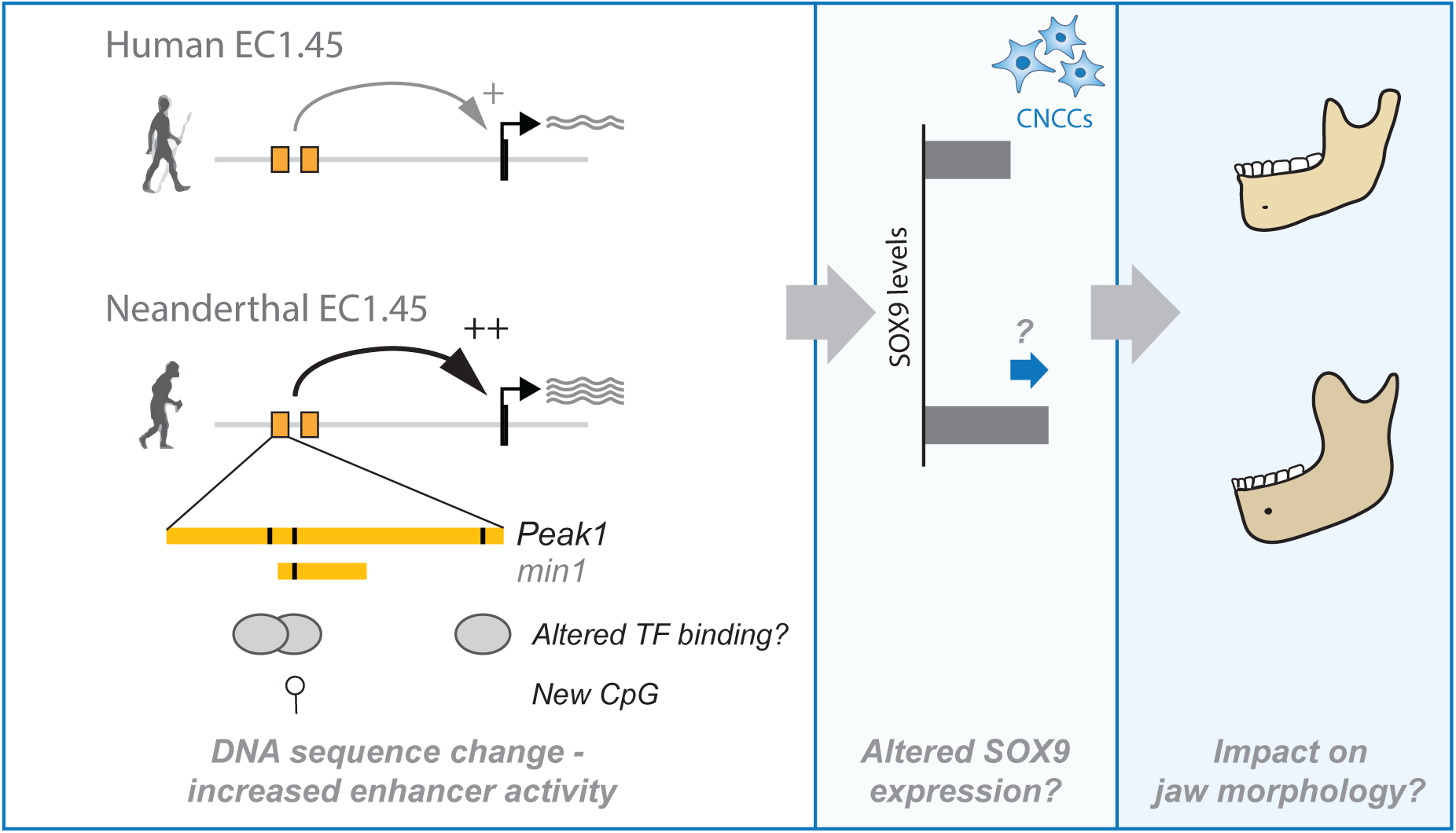
Model of increased Neanderthal enhancer activity and hypothesised impact on jaw development. A schematic illustrating the hypothesis that increased activity of Neanderthal EC1.45, perhaps driven by altered transcription factor binding or impact to CpG density and DNA methylation status, may have caused an increased level of SOX9 expression in CNCCs and their mesenchymal derivatives. This alteration may have impacted jaw formation and contributed to Neanderthal-specific alterations in jaw morphology. The potential impact of Neanderthal-derived variants on transcription factor binding is illustrated, along with the formation of a new CpG.

## Discussion

Here we have investigated the impact of three Neanderthal-derived regulatory variants on enhancer function at the *SOX9* locus in the context of craniofacial development and recent hominin evolution. Spatial expression patterns for both the human and Neanderthal enhancer during early craniofacial development appeared initially restricted to a pool of neural crest-like progenitor cells in the frontonasal region from 1 dpf, a subset of which contributed to the embryonic palate. Later, enhancer activity is also detected in a jaw-adjacent region from 2-4 dpf, between the oral ectoderm and developing Meckel’s cartilage and reaching anteriorly along the oral cavity, and at later stages appearing to extend also into the jaw joint region. These expression patterns in zebrafish embryogenesis broadly recapitulate the developmental expression patterns we previously observed in the face and limb during mouse embryogenesis, including reporter signal detected in the developing zebrafish fin (Long et al. 2020). Strikingly, although differing by only three SNVs, the Neanderthal enhancer appears to drive increased expression during early craniofacial development, especially at 2 dpf. scRNA-seq at this stage of development suggests that the two paired mandible-adjacent regions of enhancer-active cells represent mesenchymal progenitor cells of cranial neural crest origin, that are adjacent to or contribute to mature PCCs. In keeping with this, numerous EC1.45 enhancer active cells were observed to overlap with *sox10* reporter signal at 2 dpf within the presumptive PCC region of Meckel’s cartilage (Supplementary Figure 5), and by lineage-tracing we observed enhancer-active cells giving rise to chondrocytes within Meckel’s cartilage at later stages (Supplementary Figure 4). The EC1.45 enhancer cluster therefore appears to be active in a mesenchymal precartilaginous precursor pool of cells, supporting the hypothesis that perturbation of *SOX9* gene expression before cartilage specification can contribute to developmental lower jaw malformation in patients with PRS (Long et al. 2020).

During zebrafish development, CNCCs that populate the pharyngeal arches aggregate into PCCs, which are defined as regions of elevated cell density within a field of mesenchymal progenitors (Paudel et al. 2022). These PCCs will then form cartilage that defines the structure of the larval jaw (Kague et al. 2012). A number of PA1 mesenchymal and PCC markers were robustly expressed in the EC1.45 enhancer-active cell populations at 2 dpf, including *pax9*, *lhx8a* and *lhx6* which are associated with facial condensations and *barx1* which indicated contribution of cells to PCCs, as *barx1+* cells have been observed to form facial cartilages (Nichols et al. 2013; Paudel et al. 2022). It has been shown that changes to the shape and size of PCCs can translate into an altered shape of the resultant adult craniofacial structures (Paudel et al. 2022), therefore alterations to *SOX9* expression levels during development could drive changes to PCC shape, leading to structural jaw alterations (Naqvi et al. 2023).

We propose that the EC1.45 Peak1-2 enhancer-active cells adjacent to developing jaw structures from 2-4 dpf are progenitor cells expressing *sox9a*, which may contribute to forming PCCs, especially at early stages, and therefore ultimately help to shape skeletal structures of the craniofacial complex (Akiyama et al. 2005). The Neanderthal variants that we have characterised here increase the activity of EC1.45, which in the endogenous regulatory context may have driven an increase in *SOX9* expression compared to human as precartilaginous mesenchymal condensations are forming (Figure 6). While determining the implication of increased *SOX9* expression for facial development, we must consider the multiple important regulatory roles that have been demonstrated for SOX9. During facial development, SOX9 is first expressed at the neural plate border region, and in multiple species is required or sufficient for neural crest induction (Cheung and Briscoe 2003; Schock and LaBonne 2020; Spokony et al. 2002), though a proposed redundancy between SoxE factors may explain the lack of neural crest defects in other species (Akiyama et al. 2002). It has been proposed that SoxE factors may play a role in maintaining the developmental potential of neural crest cells, orthologous to SoxB1 factors during pluripotency (Buitrago-Delgado et al. 2018) and promote survival of trunk neural crest cells (Cheung et al. 2005; Sakai et al. 2006). Additionally, SOX9 plays important roles in many tissues directing progenitor cells to differentiate or proliferate in the context of both the adult organism and development, and dysregulation of SOX9 function is implicated in many cancers (Bastide et al. 2007; Jo et al. 2014). SOX9 is required for chondrogenesis and is considered the master regulator of this fate, driving upregulation of chondrocyte marker genes (Lefebvre and Dvir-Ginzberg 2017). Indeed, all osteochondrogenic cells are derived from Sox9-expressing precursors in mice (Akiyama et al. 2005) and inactivation of Sox9 in neural crest cells leads to a loss of chondrogenic potential, resulting in a switch to an osteoblast fate (Mori-Akiyama et al. 2003), with a resultant dysmorphic jaw. From our previous work, we have shown that a craniofacial SOX9 haploinsufficiency phenotype is further exacerbated by compound loss of EC1.45 from the remaining SOX9 locus (Long et al. 2020). In the context of zebrafish development, *sox9a* knockout causes craniofacial deformities (Yan et al. 2002; 2005), and drug-induced decrease in *sox9b* expression causes decreased size and number of chondrocytes resulting in jaw malformation (Xiong, Peterson, and Heideman 2008; Burns, Peterson, and Heideman 2015). We therefore hypothesise that a subtle increase of SOX9 expression in CNCCs could promote increased survival of mesenchymal progenitors or promote specification to chondrocytes. Pro-chondrogenic SOX9-target genes have been shown to be particularly sensitive to SOX9 dosage (Naqvi et al. 2023), and so very subtle changes to SOX9 expression could impact this process. In support of this, our experiments modulating levels of SOX9 expression in the zebrafish embryo revealed that overexpression of SOX9 in EC1.45-active lineages resulted in an increased size of the PCCs marked by a *sox10* reporter at 2 dpf. This is a subtle change observed in otherwise normal embryos, which could lead to changes in jaw cartilage shape, and subtly alter jaw morphology, as has been observed in the fossil record for Neanderthals compared to anatomically modern humans (Figure 6) (Bergmann et al. 2021; Rosas 2001). Future work will be needed to confirm the impact of EC1.45 Neanderthal variants at the endogenous *SOX9* locus, to quantify changes to gene expression, and further explore the consequences of increased SOX9 expression on development and ultimately jaw morphology.

Many facial features distinguishing Neanderthals from anatomically modern humans were acquired since the two lineages split from their last common ancestor (Rosas 2001; Stelzer et al. 2019). These subtle differences in lower jaw morphology between modern human and Neanderthal are hypothesised to have arisen due to alterations to mastication, different mechanical stress responses and general adaptation to environmental conditions (Bastir, O’Higgins, and Rosas 2007; Sella-Tunis et al. 2018; Stelzer et al. 2019). Interestingly, although coding sequences are generally highly conserved, genes involved in skeletal morphology have been subject to higher evolutionary change in the lineage leading to the Neanderthals than in the ancestral line common to archaic and modern human (Castellano et al. 2014). Sequence changes in the non-coding genome can drive phenotypic divergence (Frankel et al. 2011; Long, Prescott, and Wysocka 2016; Rubinstein and De Souza 2013), and have also been implicated in phenotypic differences between modern human and Neanderthal, for example in relation to skull morphology (Funato, Heliövaara, and Boeckx 2024) and vocal and facial anatomy (Gokhman et al. 2014; 2020). Here, we provide evidence that 3 Neanderthal-derived single nucleotide variants within the EC1.45 enhancer cluster may have contributed to altered craniofacial form across recent hominin evolution. Clearly, alteration of EC1.45 activity cannot account for all anatomical differences observed in the Neanderthal jaw, which was likely shaped by multiple genetic changes across multiple loci. Even at the *SOX9* locus there are multiple putative enhancer elements active during cranial neural crest development and chondrogenesis (Yao et al. 2015; Bagheri-Fam et al. 2006; Long et al. 2020; Ichiyama-Kobayashi et al. 2024) – and there can be complicated interplays between multiple enhancers acting upon a single gene (Hörnblad et al. 2021; Osterwalder et al. 2018; Bothma et al. 2015; Brosh et al. 2023; Blayney et al. 2023). That said, we understand from GWAS that craniofacial shape is a highly polygenic trait with multiple variants contributing small effects to drive morphological variation within the human population (Naqvi et al. 2022). Thus, the Neanderthal-derived SNVs within EC1.45 may contribute to the larger picture of regulatory alterations leading to mandibular morphological divergence in the Neanderthal lineage. For future work, it will be extremely valuable to understand both the mechanisms by which the EC1.45 Neanderthal SNVs result in increased activity, and what the downstream consequences of increased *SOX9* expression are during development. One or more of the Neanderthal SNVs likely alters transcription factor binding, however functional follow-up is hindered by the challenge of predicting or experimentally identifying the impacted transcription factor, especially as variants do not need to fall within a transcription factor binding site to impact factor binding or enhancer activity (Steinhaus, Robinson, and Seelow 2022; Corradin and Scacheri 2014; Claringbould and Zaugg 2021). Available tools include databases of empirically defined transcription factor binding site preference, which can help to identify candidate factors impacted by nucleotide sequence changes (Hume et al. 2015; Steinhaus, Robinson, and Seelow 2022), and machine learning methods can help to prioritise sequence variants implicated in human disease and evolutionary traits (Smith et al. 2023). Of the three SNVs, SNV2 appears to be the most promising candidate for initial investigation, as it falls within the minimally active min1 region, and generates a new CpG in the enhancer sequence, which is of added interest as EC1.45 overlaps a DMR in ancient Neanderthal bone samples compared to human (Gokhman et al. 2014; 2020; Long et al. 2020).

Together, we have demonstrated that three Neanderthal-derived SNVs in the EC1.45 enhancer cluster enhance regulatory activity and may have played a role in alteration of Neanderthal specific jaw traits through modulation of *SOX9* expression. Given the association of EC1.45 enhancer loss with human craniofacial dysmorphology, we propose that other sequence variants within this enhancer may also be associated with normal-range variation in human jaw shape (Terhune, Ritzman, and Robinson 2018), or may contribute to risk of jaw malformation upon interaction with other environmental or genetic changes. Indeed, a large number of variants can be identified across the EC1.45 enhancer cluster in the human population from the gnomAD database (Karczewski et al. 2020) and also across other primate species. Importantly, this work implicates alteration in early jaw progenitors in shaping ultimate skeletal form and highlights how regulatory function can be impacted by even very small changes to enhancer sequence. Future exploration of regulatory grammar within the EC1.45 enhancer cluster will help to uncover the impact of genetic changes on gene regulatory activity, developmental processes and resultant craniofacial morphology, relevant to both normal morphological variation and human disease.

## Materials and methods

### Zebrafish husbandry

Adult zebrafish lines were maintained as per standard protocols (Sprague et al. 2007), and carried out under a UK Home Office licence under the Animals (Scientific Procedures) Act, 1986. Zebrafish embryos were raised at 28.5°C. Embryos collected for imaging or FACS were treated with 0.003% 1-phenyl-2-thio-urea (PTU) from 24 hours post fertilisation (hpf) to prevent pigment formation.

### Generation of transgenic constructs

Dual enhancer-reporter constructs, based on the Q-STARZ system, were created by Gateway cloning (Invitrogen) as previously described (Bhatia et al. 2021; Uttley et al. 2023). The human or Neanderthal versions of EC1.45 Peak1-2 were amplified from existing plasmids by PCR using Phusion high fidelity polymerase with Gateway recombination sites included in the primer sequences (coordinates of human enhancer sequences and Neanderthal SNVs are provided in Supplementary Table 3). Enhancer sequences were then cloned into two pDONR entry vectors, pP4P1r and pP2rP3 using BP clonase. The insulator sequence was previously cloned into pDONR221 plasmid, containing 2.5 copies of the chicken HS4 sequence (Bhatia et al. 2021). The destination vector was previously synthesised by GeneArt containing a Gateway R4-R3 cassette flanked by PhiC31 and Tol2 recombination sites and minimal promoter-reporter gene units (gata2-eGFP and gata2-mCherry) (Bhatia et al. 2021). The final enhancer reporter plasmids were generated through a multi-way Gateway reaction mediated by LR clonase to combine the two enhancer sequences, the insulator, and the destination vector. Three constructs were created, NeP1P2:eGFP;HuP1P2:mCh, HuP1P2:eGFP;NeP1P2:mCh, and HuP1P2:eGFP;HuP1P2:mCh (abbreviated to Ne:GFP;Hu:Ch, Hu:GFP;Ne:Ch and Hu:GFP;Hu:Ch).

A single enhancer reporter HuEC1.45-P1P2:eGFP (abbreviated to HuP1P2:GFP) was generated similarly, as previously described (Ravi et al. 2013). The pP4P1r vector containing human Peak1-2 was combined with a pDONR221 construct containing a gata2-promoter-eGFP-polyA cassette together with a destination vector consisting of a Gateway R4-R2 cassette flanked by Tol2 recombination sites.

### Generation of stable transgenic zebrafish lines

Dual and single enhancer-reporter lines were generated as previously described (Bhatia et al. 2021; Uttley et al. 2023). Embryos were collected from wildtype AB zebrafish and injected at the one-cell stage with a 1:1 mixture of Tol2 mRNA and enhancer-reporter construct (final concentration of 25 ng/µl each). Injected F0 embryos were screened at 24 hpf for mosaic eGFP/mCherry expression and raised to adulthood. Adult F0s were crossed with wildtype AB zebrafish and the F1 progeny screened for eGFP/mCherry expression. Confirmed F0 founders were isolated with wildtype AB adults and subsequently bred for embryo collection and maintenance of the transgenic line.

### Lineage tracing in transgenic zebrafish lines

A construct to drive Cre expression under the control of the EC1.45-Peak1-2 enhancer cluster and gata2-promoter (HuP1P2:Cre) was generated by replacing eGFP from the HuP1P2:eGFP reporter plasmid with Cre amplified from a pEXPGC2(tfap2b:Cre) expression vector (Brombin et al. 2022). HuP1P2:Cre was injected at a 1:1 ratio with Tol2 mRNA (final concentration of plasmid 25 ng/µl and mRNA 35 ng/µl) into 1-cell stage embryos obtained from crossing the lineage tracing *Tg(ubi:CSY)* line with the *Tg(sox10:mRFP)* reporter line. Embryos were screened at 1 dpf using a Leica M165FCA fluorescence stereomicroscope for an expression switch from AmCyan to ZsYellow in craniofacial regions, resembling the activity pattens for the Tg(*HuP1P2:eGFP*) line.

### Transient overexpression of SOX9 during zebrafish embryogenesis

A construct to drive overexpression of human SOX9 under the control of the EC1.45-Peak1-2 enhancer cluster and gata2-promoter was generated, with T2A-eGFP included to mark injected cells. SOX9 was amplified from an Addgene plasmid (SOX9_pLX_TRC317 – TFORF2531 #142954) including a T2A sequence in the primer overhang, and sub-cloned into the HuP1P2:eGFP reporter plasmid.

Roughly 2 nL of the resultant plasmid, HuP1P2:hSOX9-T2A-eGFP, was injected into 1-cell zebrafish embryos at a 1:1 ratio with Tol2 mRNA (final concentration of plasmid 25 ng/µl, mRNA 35 ng/µl), or HuP1P2:eGFP plus Tol2 mRNA as a control. To titrate down levels of SOX9 overexpression, HuP1P2:hSOX9-T2A-eGFP and HuP1P2:eGFP plasmids were mixed 1:1 and 1:3 prior to injection, maintaining the total amount of injected DNA. Embryos were first scored at 1 dpf for phenotypic abnormalities including ‘mild’ phenotypes such as delayed growth or mild heart oedema, ‘moderate’ phenotypes indicating spinal deformities, and ‘severe’ describing embryos without observable head development. Only embryos classified as ‘normal’ without any of these phenotypes were included for imaging. Normal embryos were then screened at 1 dpf for eGFP signal using a Leica M165FCA fluorescence stereomicroscope, detection of signal in the craniofacial structures was a requirement for embryos to be included for confocal imaging and quantification.

### Live imaging

Prior to confocal imaging, embryos were treated with tricaine (MS-222) anaesthetic and screened for fluorescent signal using a Leica M165FCA or M165FC fluorescence stereomicroscope. Transgenic embryos were then mounted in 1% (w/v) low-melting point (LMP) agarose in E3 in a glass bottom dish. All images collected for mean fluorescence intensity quantification were acquired on a Nikon A1R confocal microscope using NIS elements AR software (Nikon Instruments Europe) and a 20x dry objective. A z-stack step size of 1.5µm was used, and laser power and exposure settings for each channel were the same for all images. Images of SOX9 overexpression embryos, and some enhancer-reporter embryos not used for quantification (including Supplementary Movie 1) were acquired using an Andor Dragonfly (spinning disk) confocal microscope (Andor Technologies) using Andor Fusion acquisition software. These images were captured in 40 μm pinhole mode on the iXon 888 EMCCD camera. Images were prepared for figures using FIJI (Schindelin et al. 2012) and Imaris (Oxford Instruments). 10/20x dry objectives or 40x water-immersion objectives were used for acquisition.

### Time-lapse imaging

Embryos were mounted as described above for live imaging. A portion of the LMP agarose surrounding the embryo head and body was then carefully cut away using a microsurgical knife, leaving only the tail of the embryo embedded in agarose, therefore allowing normal embryonic development throughout imaging. Embryos were covered in E3 containing MS-222 and PTU for the duration of the time-lapse imaging and maintained at 28.5°C using an Okolab bold line stage top incubator chamber. Imaging was performed as described on the Andor Dragonfly (spinning disc) confocal, acquiring a z-stack image every 60 minutes.

### Image analysis

#### Quantification of eGFP and mCherry fluorescence intensity

Enhancer activity was analysed by quantifying the mean fluorescence intensity of eGFP and mCherry in transgenic embryo images captured on the Nikon A1R. A combined eGFP/mCherry channel was created using Imaris coloc, setting the threshold for colocalization at zero. The generated colocalization channel was used to create Imaris surfaces to segment regions of signal, based on the absolute intensity thresholding of the colocalization channel with a defined surface detail of 4 µm. Segmented regions were filtered by size and classified into frontonasal or jaw regions of the face. For each surface, the mean fluorescence intensity of the original eGFP and mCherry signal was then quantified. A Wilcoxon signed-rank test was used to calculate significance p Values in R using ggPubr and plot using ggPlot2 (Wickham 2009).

#### Quantification of mRFP volumes for Tg(sox10:mRFP) embryos

The effect of SOX9 or eGFP overexpression was analysed by quantifying the volume of mRFP signal in the jaw region of injected embryos (including Meckel’s cartilage, palatoquadrate and ceratohyal regions). Using the Imaris software, a surface encompassing the entire jaw area was manually created. A blinded individual then trained an Imaris machine learning surface creation model on the mRFP signal. All images were subsequently processed using this model, thus segmenting the region of mRFP signal in the jaw in an automated manner and allowing unbiased quantification of the resulting volumes. A Bayesian mixed-effects model was used to assess differences in mRFP volume between the eGFP and SOX9 injected groups, while accounting for variability across replicates performed on different days. The model formula ‘Volume ∼ Plasmid + (1 + Plasmid | Replicate)’ was fit using the brm function from brms (Bürkner 2017). The random effects structure accounts for baseline differences in volume (intercept) and variation in the effect of plasmid (slope) between replicates. A 95% credible interval from the resulting posterior distribution was used to evaluate the magnitude and direction of any difference between groups. A Wilcoxon signed-rank test was also used to calculate significance p values using ggPubr and the data plot using ggPlot2 (Wickham 2009).

### Dissociation and sorting of 2 dpf embryos

Embryos were dechorionated and placed into 0.5x Danieau’s solution <u>(Sprague et al. 2007</u>) with tricaine (MS-222) anaesthetic. Embryo cranial regions were dissected using a scalpel blade and collected into 0.5x Danieau’s solution on ice. The samples were washed twice using 0.5x Danieau’s solution and once with FACSmax (Amsbio) after centrifugation at 300 x g for 1 min at 4°C. Finally, samples were resuspended in 500 μL FACSmax and passed through a 35-μm cell strainer to obtain single-cells. Cells were sorted based on fluorescence using the CytoFLEX SRT (Beckman Coulter).

### Single cell RNA-sequencing

Up to 40,000 sorted cells per sample were processed for scRNA-seq and Illumina library preparation using the 10x Genomics Chromium single-cell 3′ gene expression technology (v3.1) (Zheng et al. 2017) by the IGC FACS facility. Sequencing of the resulting libraries was carried out on the NextSeq 2000 platform (Illumina Inc.) using the NextSeq 1000/2000 P3 Reagents (100 cycles) v3 Kit, aiming for ∼50,000 reads per cell. Cell Ranger (10x Genomics, v7.1.0) was used to create FASTQ files and perform alignment, filtering, barcode and UMI counting. A custom reference genome was created from GRCz11 using the Lawson Lab transcriptome annotation (downloaded from https://www.umassmed.edu/lawson-lab/reagents/zebrafish-transcriptome/, (Lawson et al. 2020)) and manually annotated eGFP and mCherry sequences.

### Analysis of single cell RNA-seq

#### Cell calling and quality control

Cell calling was performed using the emptyDrops function from DropletUtils, excluding mitochondrial and ribosomal genes to improve the filtering of barcodes corresponding to ambient RNA or cell fragments (Lun et al. 2019). QC was then performed using the scater package to remove cells with detected genes < 400, library size <1000, and mitochondrial reads >5% (McCarthy et al. 2017). Further QC was carried out using SoupX to identify and remove cells with high levels of ambient RNA (Young and Behjati 2020). Cell counts after each stage of filtering are shown in Supplementary Table 4.

#### Clustering and cell type identification

Clustering was performed using Seurat (v5, (Hao et al. 2024)). Log normalisation, scaling, and highly variable gene (HVG) detection was carried out using SCTransform, with regression of mitochondrial expression and cell-cycle stage. Seurat CellCycleScoring was used to assign cell cycle stages. Principle component analysis was performed using the HVG list, which were then used for K-nearest neighbour analysis and Louvain clustering. To identify cell types, Seurat FindAllMarkers was used to identify cluster gene markers, and the Daniocell resource (https://daniocell.nichd.nih.gov/index.html, (Sur et al. 2023)) was used as a reference for cell type mapping using SingleR (Aran et al. 2019). At this stage, a small number of clusters corresponding to contaminating cell types were removed (blood, skin, and muscle tissues) to improve clustering resolution for the cell types of interest (see Supplementary Table 4). Final clustering was carried out using the following parameters - sample *Tg(Ne:GFP;Hu:Ch)* – 10 dimensions, resolution 0.5; sample *Tg(Hu:GFP;Ne:Ch)* – 15 dimensions, resolution 0.2.

#### Quantification of eGFP and mCherry reads

Cells with at least 3 eGFP/mCherry reads were assigned as enhancer-active (human, Neanderthal, or double-positive).

**Supplementary Figure 1.**

(A) Representative image of the cranial region of embryos at 1 dpf for *Tg(HuEC1.45-P1P2:eGFP)* crossed with *Tg(sox10:mRFP)*. Maximum intensity projections from confocal microscopy. Scale bar 100 µm.

(B) Schematics of zebrafish embryonic cranial region (ventral and lateral views) indicating the location of HuEC1.45-P1P2 enhancer activity in relation to the *col2a1a:RFP* reporter signal in precartilaginous condensates (PCCs) which mature to form cartilage templates from 2-4 dpf. m – Meckel’s; p – palatoquadrate; e – ethmoid plate; c – ceratohyal.

(C) Representative images of the cranial region of embryos at 2, 3 and 4 dpf for *Tg(HuEC1.45- P1P2:eGFP)* crossed with *Tg(col2a1a:RFP)*. Maximum intensity projections from confocal microscopy. Scale bars 100 µm.

(D) Representative images of the pectoral fin for embryos at 3 and 4 dpf for *Tg(HuEC1.45- P1P2:eGFP)* crossed with *Tg(col2a1a:RFP)*. Maximum intensity projections from confocal microscopy. Scale bars 100 µm.

**Supplementary Figure 2.**

(A) Representative confocal images (maximum intensity projections) for embryos at 1 dpf for the *Tg(Ne:GFP;Hu:Ch)*, *Tg(Hu:GFP;Ne:Ch)* and *Tg(Hu:GFP;Hu:Ch)* lines. Ventral view of cranial region. Scale bars 100 µm.

(B) Representative confocal images (maximum intensity projections) for embryos at 3 dpf for the *Tg(Ne:GFP;Hu:Ch)*, *Tg(Hu:GFP;Ne:Ch)* and *Tg(Hu:GFP;Hu:Ch)* lines. Ventral view of cranial region. Scale bars 100 µm.

(C) Illustrative images of surfaces used to quantify mean fluorescence intensity of eGFP and mCherry signal for 1, 2 and 3 dpf. At 1 dpf, the surface includes the entire domain of signal. At 2- 3 dpf, the surface is separated into a frontonasal region, including some cells contributing to the developing palate (pink), and the mandible-adjacent region (blue).

(D) Quantification from (A) at 1 dpf for the entire segmented region. P values from a Wilcoxon signed-rank test are shown.

(E) Quantification from (B) at 3 dpf for the jaw region (upper) and frontonasal region (lower). P values from a Wilcoxon signed-rank test are shown.

**Supplementary Figure 3.**

(A) FACS plots for cells dissected from the cranial region of *Tg(Hu:GFP;Ne:Ch)* and *Tg(Ne:GFP;Hu:Ch)* transgenic lines highlighting cells that were collected for 10X scRNA-seq (pink gate, approximate gating used for sorting). Matched non-fluorescent cells from wildtype embryo cranial regions were used for setting the sorting gates.

(B) Dot plot illustrating average expression level and percentage of cells expressing markers for each cluster shown in Figure 4C.

(C) Dot plot illustrating CNCC, frontonasal, pharyngeal arch 1 and mesenchymal condensation marker genes.

(D) Expression of *eGFP*, *mCherry*, *sox9a* and *sox9b* in single cells visualised on UMAP plots, as for Figure 4C.

(E) Cells from *Tg(Ne:GFP;Hu:Ch)* embryos at 2 dpf, visualised on a UMAP plot. Cell types were annotated to clusters aided by marker gene expression and the Daniocell atlas.

(F) Dot plot showing average expression level and percentage of *Tg(Ne:GFP;Hu:Ch)* cells expressing markers for each cluster shown in (E).

(G) Dot plot showing average expression level and percentage of *Tg(Ne:GFP;Hu:Ch)* cells expressing eGFP, mCherry, *sox9a* or *sox9b* (clusters from E).

**Supplementary Figure 4.**

(A) Schematic of a cross between the *Tg(ubi:CSY)* reporter line (full name *Tg(ubi:LoxP-AmCyan-LoxP-ZsYellow)*) and *Tg(sox10:mRFP)*. Resultant embryos were injected with the HuP1P2:Cre construct along with Tol2 mRNA.

(B) (i) Confocal images (maximum intensity projections) for representative embryo 1, for which cells in the frontonasal craniofacial region at 1 dpf exhibited a Cre-mediated AmCyan to ZsYellow switch (arrowheads, lateral images). At 2 dpf, ZsYellow signal is observed adjacent to the Meckel’s precartilaginous condensate (PCC) (arrow). Across 12 hours of time-lapse imaging from 2 dpf, a ZsYellow-positive, sox10-positive cell is observed to appear overlapping the developing Meckel’s PCC region, a white arrow highlights an outlined cell for reference which is proximal to the emerging ZsYellow-positive cell, marked by an arrowhead (see also Supplementary movie 6). M – Meckel’s PCC.

(A) (ii) Confocal images (maximum intensity projections) for representative embryo 2, for which ZsYellow-positive cells were identified at 3 dpf adjacent to the Meckel’s cartilage, proximal to ZsYellow-positive, sox10-positive Meckel’s chondrocytes (white arrowheads). At 4 dpf, elongation of the ZsYellow-positive chondrocytes can be observed (white arrowheads).

(B) (iii) Confocal images (maximum intensity projections) for representative embryo 3, for which ZsYellow-positive, sox10-positive cells were identified at 3 dpf adjacent to and within the Meckel’s cartilage (white arrowhead). Scale bars 20 µm. Objective used for imaging is indicated in upper left corner of images. M – Meckel’s.

**Supplementary Figure 5.**

(A) Representative confocal microscopy image (maximum intensity projection) showing a ventral view of the embryonic cranial region for a 2 dpf embryo from a cross between the *Tg(HuEC1.45- P1P2:eGFP)* and *Tg(sox10:mRFP)* lines. eGFP-positive, sox10-positive cells in Meckel’s precartilaginous condensate are indicated by white arrowheads (double-positive cells within Meckel’s PCC were identified for 4/4 embryos imaged).

(B) As for (A), showing a ventral view of a 2 dpf embryo from a cross between the *Tg(HuEC1.45- P1P2:eGFP)* and *Tg(col2a1a:RFP)* lines. eGFP-positive, sox10-positive cells in Meckel’s precartilaginous condensate are indicated by white arrowheads.

(C) Representative confocal microscopy image (maximum intensity projections) showing a ventral view of the embryonic cranial region for a 2 dpf embryo from a cross between the *Tg(Ne:GFP;Hu:Ch)* and *Tg(sox10:mRFP)* lines. eGFP-positive, sox10-positive cells in Meckel’s precartilaginous condensate are indicated by white arrowheads. Scale bars 50 µm.

**Supplementary Figure 6.**

(A) Quantification of phenotypes observed at 1 dpf after injection at the 1-cell stage with either HuEC1.45-P1P2:SOX9-T2A-eGFP or HuEC1.45-P1P2:eGFP. Mild phenotypes include common abnormalities such as delayed growth or mild heart oedema, moderate phenotype indicates spinal deformities, while severe denotes embryos without observable cranial development.

(B) Images of *Tg(sox10:mRFP)* embryos injected with either HuEC1.45-P1P2:eGFP or HuEC1.45-P1P2:SOX9-T2A-eGFP at 1 dpf. eGFP expression can be seen in the developing craniofacial region (black arrowheads).

(C) Representative images of segmented regions of *sox10:mRFP* expressing cells in the developing jaw and oral-adjacent regions for either HuEC1.45-P1P2:eGFP or HuEC1.45- P1P2:SOX9-T2A-eGFP injected embryos. The *Tg(sox10:mRFP)* reporter was used to create a surface encompassing the mesenchymal cells of Meckel’s and palatoquadrate precartilaginous condensations and the regions extending along the oral ectoderm. Scale bars 50 µm.

**Supplementary Movie 1.**

Animated movie of a confocal z-stack in 3D illustrating the location of eGFP-positive cells in relation to mRFP in the developing jaw precartilaginous condensations at 2 dpf for a cross between the transgenic lines *Tg(HuP1P2:GFP)* and *Tg(sox10:mRFP)*. Prepared in Imaris, scale bar is indicated.

**Supplementary Movie 2.**

Animated movie of a confocal z-stack in 3D showing eGFP-positive and mRFP-positive cells in the developing jaw region as surfaces at 2 dpf for a cross between the transgenic lines *Tg(HuP1P2:GFP)* and *Tg(sox10:mRFP)*. eGFP-positive regions form paired structures at the hinge region adjacent to the oral cavity, and proximal to the forming Meckel’s precartilaginous condensation. The sox10:mRFP surface encompasses the mesenchymal cells of Meckel’s and palatoquadrate precartilaginous condensations and mesenchymal cells extending along either side of the oral ectoderm. Prepared in Imaris, scale bar is indicated.

**Supplementary Movie 3.**

Animated movie of a confocal z-stack in 3D showing eGFP-positive and mRFP-positive cells in the developing jaw region as surfaces at 4 dpf for a cross between the transgenic lines *Tg(HuP1P2:GFP)* and *Tg(sox10:mRFP)*. eGFP-positive cells form paired regions adjacent and lateral to the Meckel’s cartilage, with signal extending into and around the jaw joint between Meckel’s and the palatoquadrate cartilage template (marked by sox10:mRFP). Prepared in Imaris, scale bar is indicated.

**Supplementary Movie 4.**

Time-lapse movie of confocal images (maximum intensity projections) from around 2-4 dpf for an embryo from a cross between transgenic lines T*g(HuP1P2:GFP)* and *Tg(sox10:mRFP)*. Scale bar 100 µm.

**Supplementary Movie 5.**

Time-lapse movie of confocal images (maximum intensity projections) from Supplementary Movie 4, cropped and focused on the developing embryonic palate region. White arrow indicates eGFP-positive cells in the ethmoid plate (embryonic palate). Scale bar 20 µm.

**Supplementary Movie 6.**

Time-lapse movie of confocal images (maximum intensity projections) over 12 hours from 2 dpf of representative embryo 1 (see Supplementary Figure 4Bi), which exhibited an AmCyan to ZsYellow switch in the frontonasal craniofacial region at 1 dpf. ZsYellow signal was observed adjacent to Meckel’s cartilage at 2 dpf, with emergence of a ZsYellow-positive, sox10-positive cell within Meckel’s cartilage observed at around 60 hpf (white arrow). Scale bar 20 µm.

**Supplementary Table 1**

Expression of cluster marker genes in scRNA-seq data, called using Seurat FindAllMarkers.

**Supplementary Table 2.**

Expression of eGFP (≥3 reads, green), mCherry (≥3 reads, magenta) or both (≥3 reads in total for eGFP and mCherry, yellow) across scRNA-seq clusters.

**Supplementary Table 3.**

Genomic coordinates for human EC1.45 and Neanderthal SNVs (hg19).

**Supplementary Table 4.**

Cell counts after each stage of quality control and filtering of scRNA-seq datasets.

## Acknowledgements

We thank Joanna Wysocka for advice and support at the conception of this project. We are grateful to Cameron Wyatt and the Zebrafish facility at the Institute of Genetics and Cancer (IGC), Ann Wheeler and the Advanced Imaging Resource at the IGC, and the Flow Cytometry facility at the IGC for their technical support and advice with experiments, and the Wellcome Trust Clinical Research Facility for sequencing support. We are grateful to Richard White for sharing the *Tg(ubi:LoxP-AmCyan-LoxP-ZsYellow)* zebrafish line. We thank Shipra Bhatia and Wendy Bickmore for advice with the Q-STARZ assay, Laura Murphy and James Iremonger for help with image analysis, Andrew Badrock, Erika Kague, and Jana Travnickova for advice with zebrafish embryo phenotyping, Charli Corcoran for assistance screening zebrafish founders, Elizabeth Freyer for FACS analysis, Susan Campbell for Chromium 10X sample preparation, and Hywel Dunn-Davies for support with statistics. We are also grateful to Wendy Bickmore for feedback on the manuscript. K.U. was funded by an MRC Transition fellowship (WT13290025), H.J.J. was funded by ERASMUS+, E.O. and H.J.J. are funded by an MRC PhD Studentship (MC_ST_00035) and H.K.L. is supported by an MRC University Unit grant (MC_UU_00035/12).

