## Supplementary Figures 1-6 for "Neanderthal-derived variants shape craniofacial enhancer activity at a human disease locus"

Supplementary Figure 1

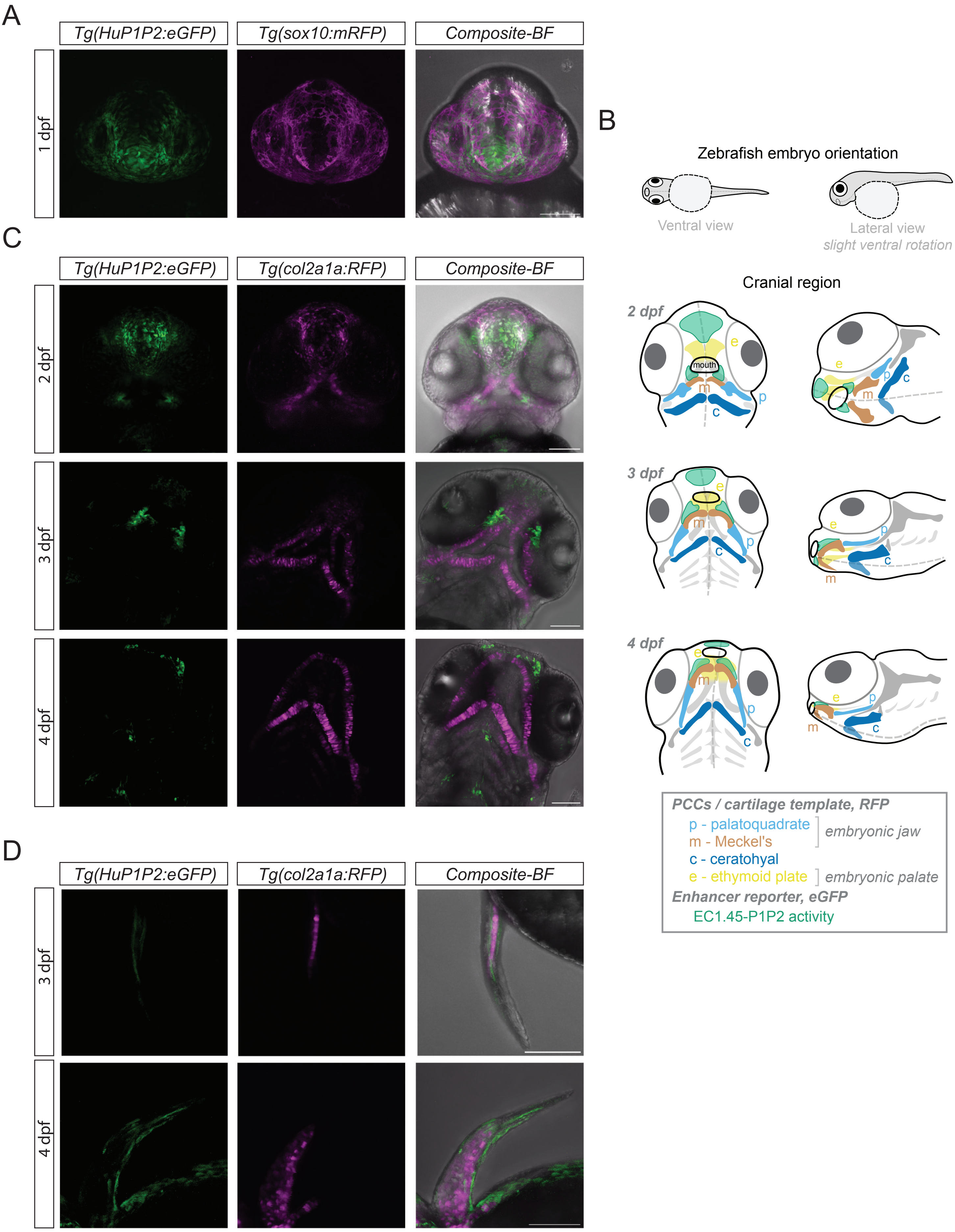

Supplementary Figure 2

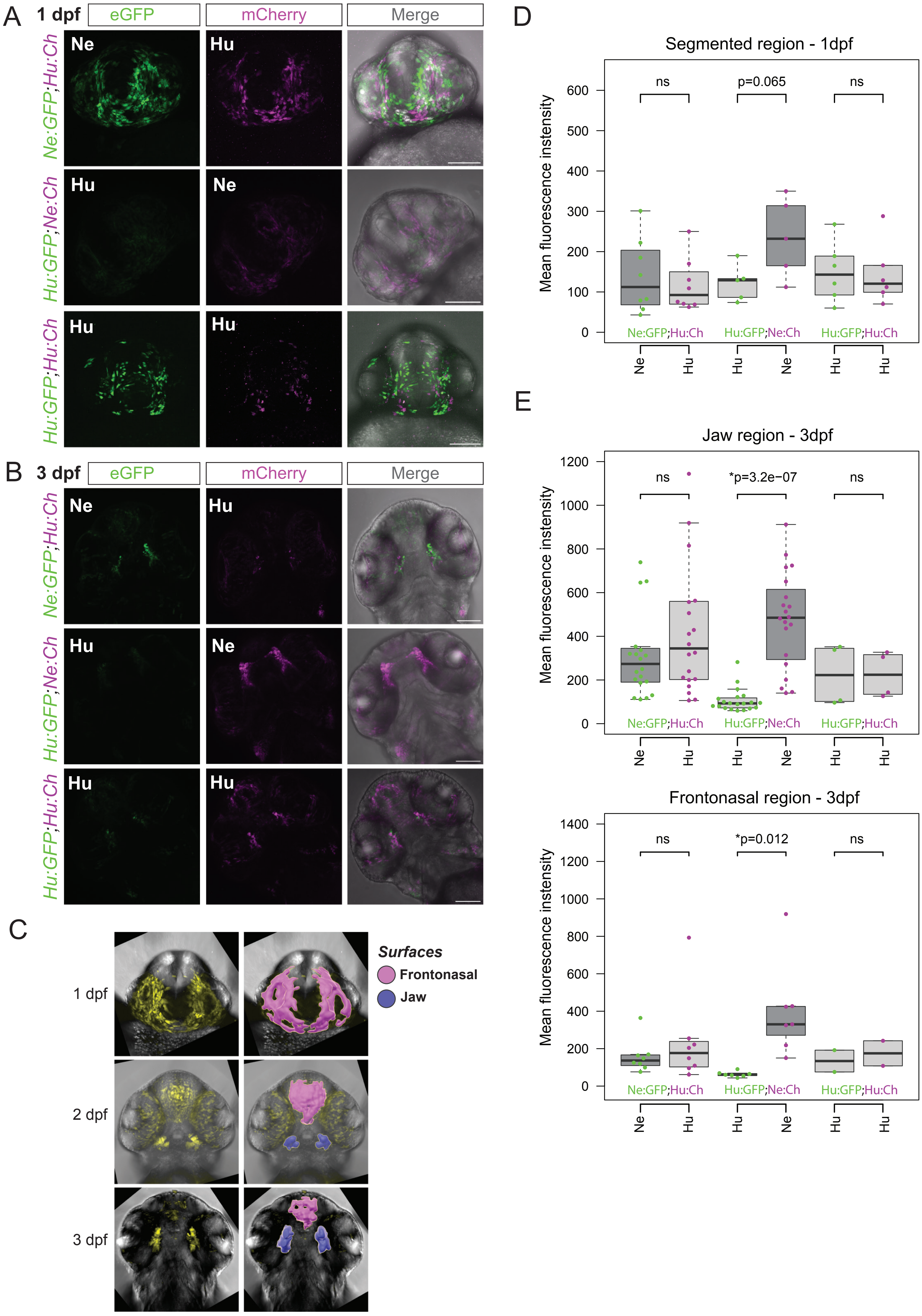

### Supplementary Figure 3

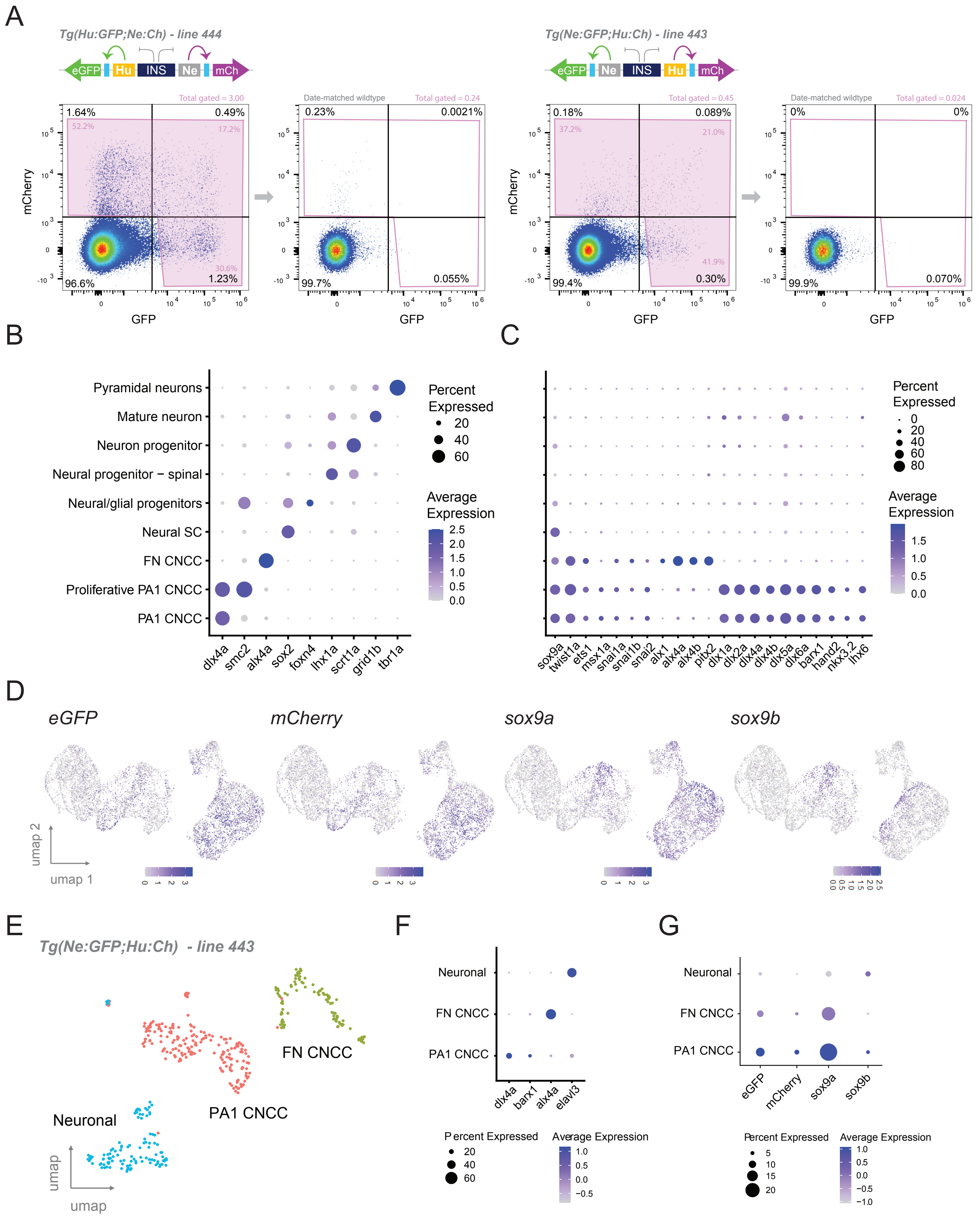

Supplementary Figure 4

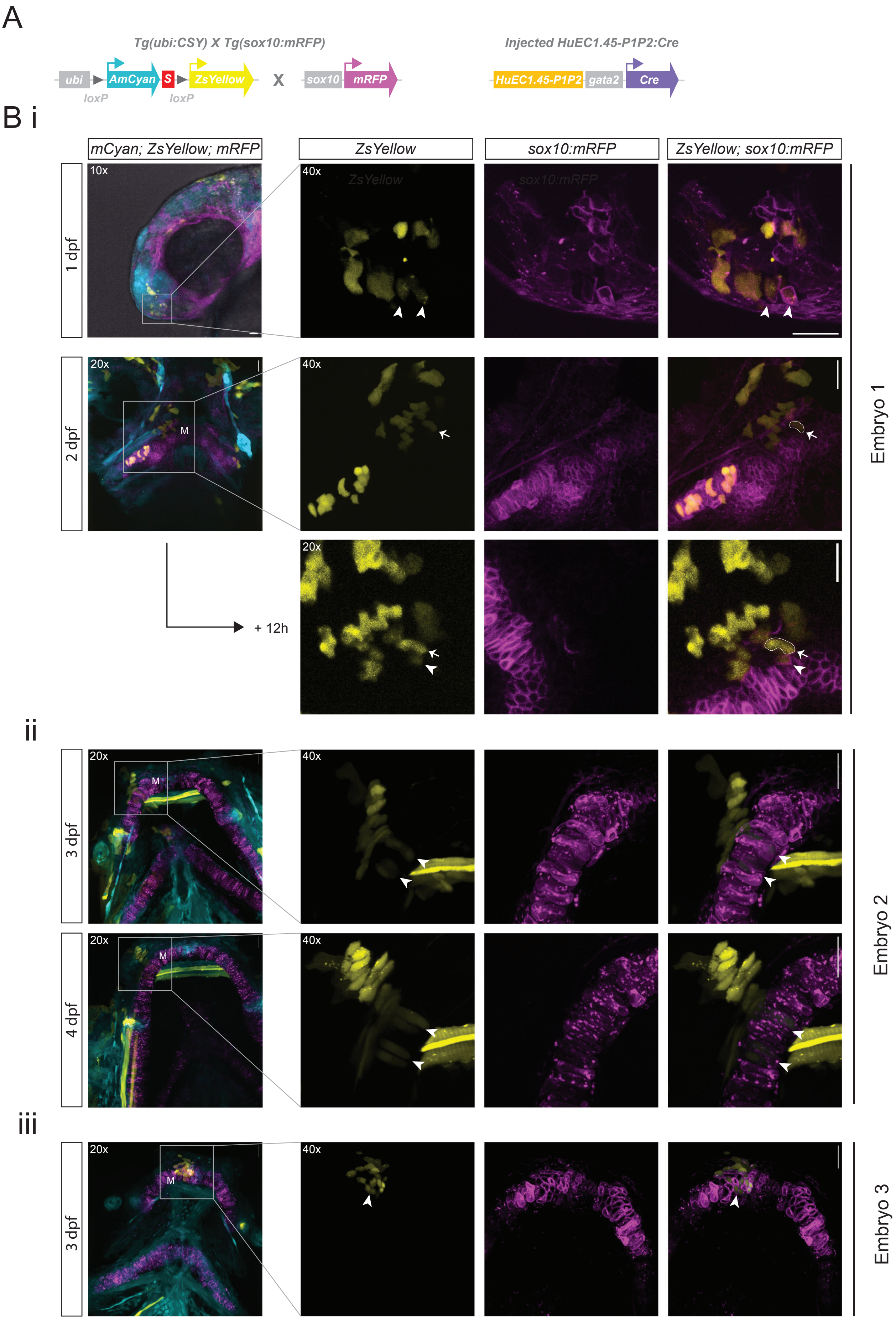

Supplementary Figure 5

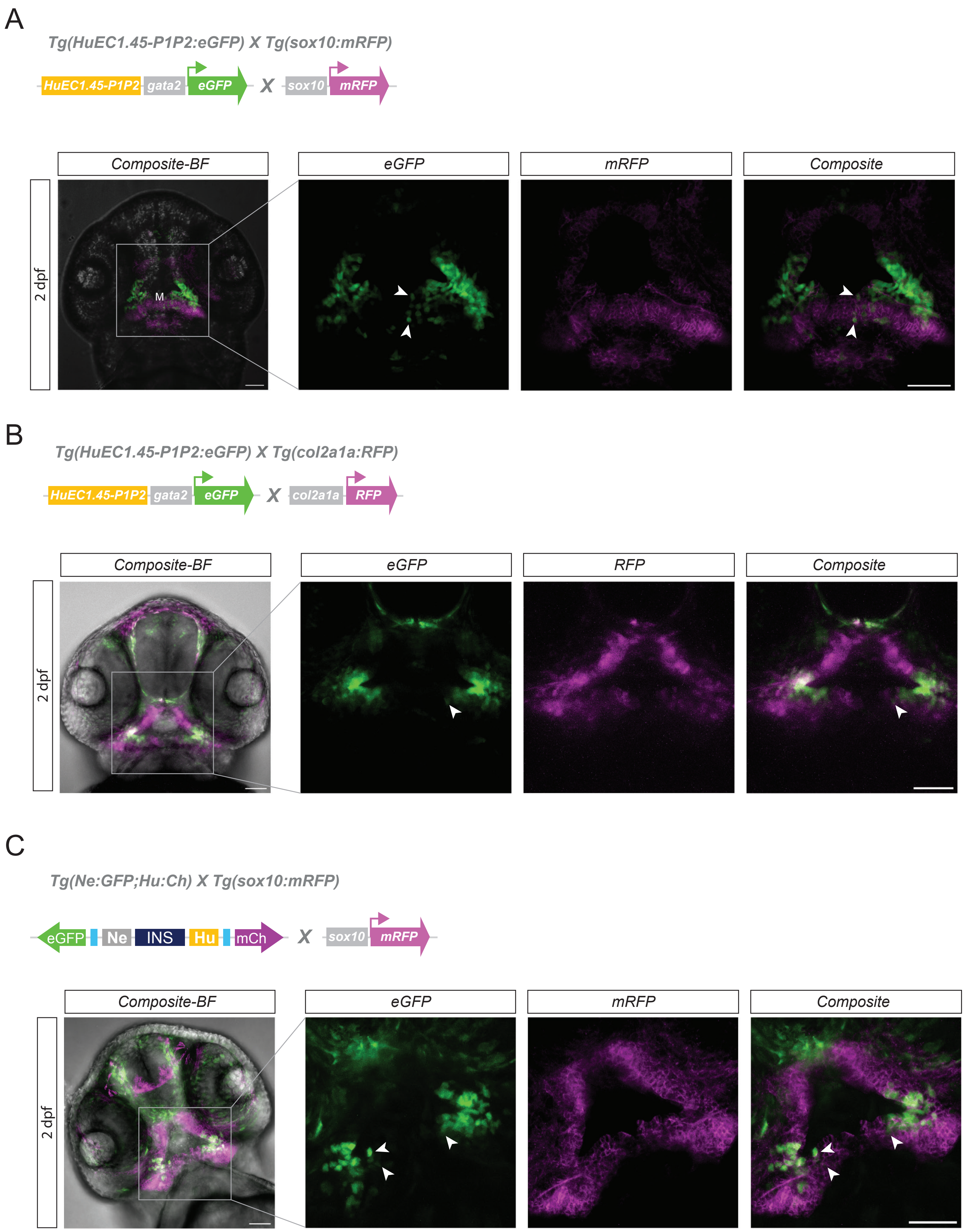

Supplementary Figure 6

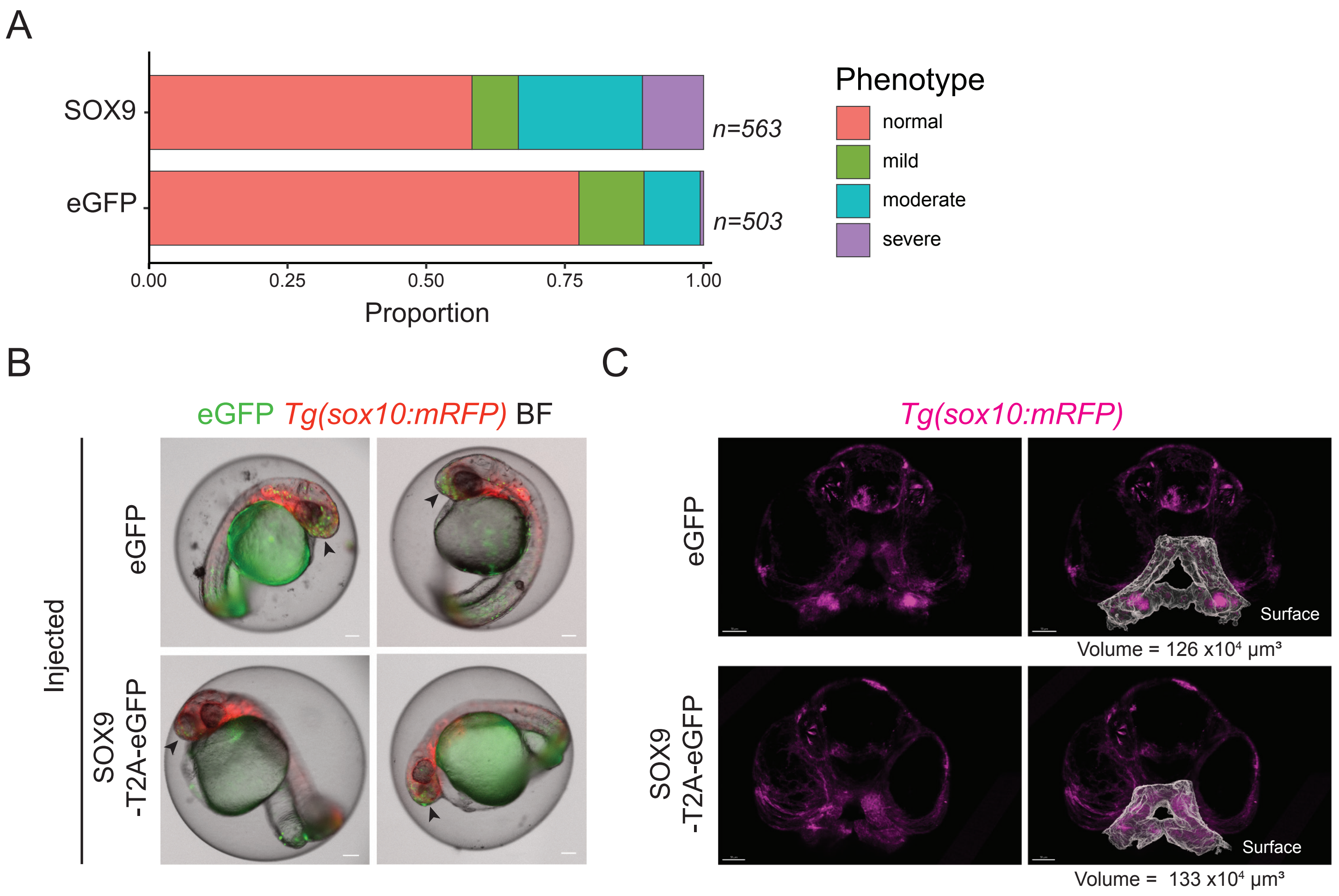
